# Parallel evolution of multiple mechanisms for demethylase inhibitor fungicide resistance in the barley pathogen *Pyrenophora teres* f. sp. *maculata*

**DOI:** 10.1101/798991

**Authors:** Wesley J. Mair, Geoffrey J. Thomas, Kejal Dodhia, Andrea L. Hills, Kithsiri W. Jayasena, Simon R. Ellwood, Richard P. Oliver, Francisco J. Lopez-Ruiz

## Abstract

The demethylase inhibitor (DMI) or group 3 fungicides are the most important class of compounds for the control both of plant and human fungal pathogens. The necrotrophic fungal pathogen *Pyrenophora teres* f. sp. *maculata* (*Ptm*), responsible for spot form of net blotch (SFNB), is currently the most significant disease of barley in Australia, and a disease of increasing concern worldwide. The main basis for management of SFNB is by fungicide application, and in Australia the DMIs predominate. Although reduced sensitivity to DMI fungicides has recently been described in the closely related pathogen *P. teres* f. sp. *teres* (*Ptt*), the mechanisms of DMI resistance have not thus far been described for *Ptm*. In this study, several different levels of sensitivity to DMI fungicides were identified in Western Australian strains of *Ptm* from 2016 onwards, and reduced sensitivity phenotypes were correlated with a number of distinct mutations in both the promoter region and coding sequence of the DMI target gene encoding cytochrome P450 sterol 14α-demethylase (*Cyp51A*). Five insertions elements of 134-base pairs in length were found at different positions within the upstream regulatory region of *Cyp51A* in both highly DMI-resistant (HR) and select moderately DMI-resistant (MR1) *Ptm* isolates. The five insertion elements had at least 95% sequence identity and were determined to be Solo-LTR (Long Terminal Repeat) elements, all deriving from *Ty1*/*Copia*-family LTR Retrotransposons. The 134-bp elements contained a predicted promoter sequence and several predicted transcription factor binding sites, and the presence of an insertion element was correlated with constitutive overexpression of *Cyp51A*. The substitution of phenylalanine by leucine at position 489 of the predicted amino acid sequence of CYP51A was found in both HR and select moderately DMI-resistant (MR2) *Ptm* isolates. The same F489L amino acid substitution has been previously reported in Western Australian strains of *Ptt*, where it has also been associated with reduced sensitivity to DMI fungicides. In *Ptm*, the F489L amino acid change was associated with either of three different single nucleotide polymorphisms in codon 489. This suggests that, in contrast to *Ptt*, in *Ptm* the F489L mutation has emerged as a result of three distinct mutational events. Moderately DMI-resistant isolates had one or the other of the F489L substitution or a promoter insertion mutation, whereas highly DMI-resistant isolates were found to have combinations of both mechanisms together. Therefore, multiple mechanisms acting both alone and in concert were found to contribute to the observed phenomena of DMI fungicide resistance in *Ptm*. Moreover, these mutations have apparently emerged repeatedly and independently in Western Australian *Ptm* populations, by a process of convergent or parallel evolution.

## INTRODUCTION

The demethylase inhibitor (DMI) or group 3 fungicides are the most important class of compounds for the control both of plant and human fungal pathogens (Cools, Hawkins, and Fraaije 2013). DMIs are a structurally diverse class of compounds (Chen et al. 2014), and act by inhibiting the activity of the enzyme lanosterol 14α-demethylase cytochrome P450 monooxygenase (CYP51), which is involved in the pathway of ergosterol biosynthesis (Ziogas and Malandrakis 2015). Ergosterol is the main sterol of the cell membrane in most fungi and is essential for maintaining cell membrane integrity and permeability (Rehfus 2018). The CYP51 enzyme catalyses the removal by oxidation of the methyl group of lanosterol at C14 (Mullins et al. 2011). The DMI compound acts as a non-competitive inhibitor by binding with a nitrogen atom of the heterocyclic ring to the haem group iron within the active site of the enzyme, preventing the transfer of O_2_ to the substrate (Mullins et al. 2011); while the side chains of the DMI compound interact with residues within the catalytic pocket, and this interaction determines the binding affinity (and therefore the inhibitory effect) of a particular DMI compound to a given CYP51 (Warrilow et al. 2013). The result of this is a depletion of ergosterol, an accumulation of 14α-methylated precursor sterols, and lethal membrane disruption (Ziogas and Malandrakis 2015, Becher and Wirsel 2012).

Resistance to DMI fungicides is an increasing concern in both the field and the clinic (Ribas E Ribas et al. 2016). Four main mechanisms have been observed for resistance to DMI fungicides in filamentous fungi (Ziogas and Malandrakis 2015): i) Increased efflux. Overexpression of genes encoding membrane-bound drug transporters, such as those of the major facilitator superfamily or ATP-binding cassette transporters, leads to reduced accumulation of the fungicide within the cell (Ziogas and Malandrakis 2015). Due to the limited substrate specificity of these transporters, overexpression generally confers multi-drug resistance, that is resistance across multiple unrelated classes of fungicides (Cools, Hawkins, and Fraaije 2013). Enhance efflux by overexpression of transporters is the mechanism most frequently implicated in DMI resistance in fungal pathogens of humans (Ziogas and Malandrakis 2015), but among plant pathogenic fungi this mechanism has been observed only in comparatively few species such as *Botrytis cinerea* (Kretschmer et al. 2009) and *Penicillium digitatum* (Nakaune et al. 1998). ii) Overexpression of CYP51. Increased production of the DMI target is a common mechanism of resistance in pathogenic fungi of plants (Cools, Hawkins, and Fraaije 2013) and has also been observe in human pathogens (Becher and Wirsel 2012). Most frequently the overexpression is constitutive (Cools, Hawkins, and Fraaije 2013) and is generally linked to alterations in the regulatory region upstream of the target gene *Cyp51* (Ziogas and Malandrakis 2015). These changes are most often insertion sequences, with powerful promoters, and which bear homology to transposable elements (TE) (Cools, Hawkins, and Fraaije 2013). Target overexpression generally confers a level of resistance across multiple DMI compounds, sometimes conferring complete DMI cross-resistance (Cools, Hawkins, and Fraaije 2013); however the levels of resistance are in most cases relatively lower than those conferred by target site alteration (Ziogas and Malandrakis 2015). iii) Target site modification of CYP51. Changes in the polypeptide structure of CYP51 resulting in a reduced binding affinity to DMIs has been described in many species (Ziogas and Malandrakis 2015). Alterations of the residues interacting with the side-chains of the structurally heterogeneous DMIs means that target site modification typically results in a differential pattern of sensitivity across various DMI compounds, and incomplete cross-resistance, with a particular mutation most affecting sensitivity to a specific compound or group of structurally similar compounds (Cools, Hawkins, and Fraaije 2013), and in some cases negative cross-resistance is reported (Fraaije et al. 2007). iv) Multiple CYP51s. The existence of multiple paralogs of CYP51 has been reported in several species (Becher and Wirsel 2012). By convention, the paralog carried by all ascomycetes is termed CYP51B, while divers species also retain a second paralog termed CYP51A, the result of an ancient gene duplication event (Hawkins et al. 2014). *Fusarium* species carry a third unique paralog termed CYP51C, while in other species carrying a third CYP51 it is the result of more recent gene duplications of either the CYP51A or CYP51B paralog (Becher and Wirsel 2012). In those species carrying multiple paralogs it is most frequently the CYP51A that is mutated, overexpressed or otherwise implicated in DMI resistance (Fan et al. 2014, Cools, Hawkins, and Fraaije 2013, Ziogas and Malandrakis 2015). The presence of multiple CYP51 paralogs may result in an inherent reduced sensitivity to DMIs (Cools, Hawkins, and Fraaije 2013), or the retention of multiple, functionally-redundant gene copies may mitigate any biological costs of fungicide resistance mutations (Ziogas and Malandrakis 2015). A combination of multiple mechanisms together has also been reported, for instance in *Aspergillus fumigatus* (Mellado et al. 2007, Snelders et al. 2015), *Pyrenopeziza brassicae* (Carter et al. 2014), and *Zymoseptoria tritici* (Cools et al. 2012), the combination of target site alterations with insertions in the promoter leading to CYP51 overexpression was associated with the highest observed levels of DMI resistance and cross-resistance against all DMIs tested.

Recently, reduced sensitivity to DMI fungicides has been described in *Pyrenophora teres* Drechsler f. sp. *teres* [Sacc.] Shoemaker (*Ptt*), responsible for net form of net blotch (NFNB) of barley (*Hordeum vulgare*), and the molecular bases of this reduction in sensitivity characterised (Mair, Deng, et al. 2016). From 2013 onwards, strains were identified showing resistance factors ranging from 1.1 for epoxiconazole to 31.7 for prochloraz, and this resistant phenotype was correlated with the presence of a substitution of phenylalanine by leucine at position 489 in one of the CYP51A paralogs. This F489L mutation, which was found to be homologous to the mutations F495I in the CYP51A of *A. fumigatus* and to F506I in the CYP51B of *Penicillium digitatum*, produced a substantial conformational rearrangement within the heme cavity, resulting in the constriction of regions adjacent to the docking sites of the DMIs tebuconazole, difenoconazole and prochloraz, and lower predicted binding affinities for these fungicides. The *Cyp51A* genes (two paralogous copies identified) were not found to be expressed constitutively in either the sensitive or resistant *Ptt* strains, but inducible overexpression of all the *Cyp51* genes was observed in resistant strains with the addition of the DMI compound tebuconazole. This phenotype could not be linked to any mutations in either of the *Cyp51A* or *Cyp51B* promoter regions, nor were any mutations identified in the coding sequence of the paralog *Cyp51B*.

The necrotrophic fungal pathogen *Pyrenophora teres* f. sp. *maculata* Smedegard-Petersen (*Ptm*), the closely related *forma specialis* responsible for spot form of net blotch (SFNB) of barley, is a disease of increasing concern worldwide (Burlakoti et al. 2016). SFNB is currently the most significant disease of barley in Australia, causing an average of AUD$43 million in losses annually, with potential losses estimated at AUD$192 million annually in the absence of current control measures (Murray and Brennan 2010). Owing to a lack of highly resistant cultivars (Trainor 2018), the main basis for management of SFNB is by fungicide application (Sierotzki et al. 2007, Akhavan et al. 2017), with the DMIs the most important class of compounds used to control this disease in Australia (APVMA 2019). While there are currently some reports regarding reduced sensitivity to DMI fungicides in SFNB (Akhavan et al. 2017, Campbell and Crous 2002), the molecular bases underlying DMI resistance in *Ptm* have not thus far been addressed. In this study, several different levels of sensitivity to DMI fungicides were identified in Western Australian strains of *Ptm* from 2016 onwards, and reduced sensitivity phenotypes were correlated with a number of distinct mutations in both the promoter and coding sequences of *Cyp51A*.

## MATERIALS & METHODS

### Fungal isolates

Samples of SFNB diseased barley were collected by a combination of especially designed bait trials, field trips and a network of collaborators. Bait trials located in Inverleigh (Victoria), and in Esperance, Northam, and Mt Barker (Western Australia), were sown with the SFNB susceptible varieties SY Rattler, Fathom, Spartacus, Stirling and Planet, and were designed to work as a fungicide resistance catch system. Plots of 2m × 4m were sprayed with either 1× or 2× the maximum registered dose of fungicides from the Groups 3 (DMI), 11 (QoI) and 7 (SDHI), at growth stages GS31 and GS39. Treatments were replicated three times. Leaf samples from bait trials were collected seven days following the second spray application. Leaves were air dried for at least 24 h and subsequently stored at room temperature.

Small tissue fragments containing lesions were cut from dried leaf samples and disinfected for 15 s in 70% (v/v) ethanol, 30 s in 1% (v/v) NaOCl, and washed for 60 s in sterile deionized H_2_O. Surface-sterilized lesions were air dried for 30 min and then inoculated into 1.5% (w/v) agar plates amended with the antibiotics ampicillin (100 μg mL^−1^), streptomycin (30 μg mL^−1^), and neomycin (50 μg mL^−1^). After incubation at room temperature for 7 days, colony margins were sub-cultured to V8-potato-dextrose agar (10 g potato-dextrose agar, 3 g CaCO3, 15 g agar, 150 mL V8 juice in 850 mL deionized H_2_O; V8PDA) plates. For the induction of sporulation, hyphae of 5 day cultures were flattened gently with sterile deionized H_2_O using a glass spreader; the plate was then incubated at room temperature under near-UV and white light for 16 h, followed by 15 °C in darkness for 24 h. A single conidium was transferred to a new V8PDA plate amended with the three antibiotics at the concentrations described above; 4 mm-diameter mycelial plugs taken from the colony edge of this culture were stored at −80 °C and used for all subsequent testing. Isolates characterised in this study are listed in Table 1, a complete list of isolates screened in the course of this study is included in Supplementary Materials Table S1.

**Table 1.** Isolates of *Pyrenophora teres* f. sp. *maculata* characterised in this study.

| Sampling year | Location | Phenotype | Isolate(s) |
| --- | --- | --- | --- |
| 1996 | Badgingarra, WA | S | SG1 |
| 2001 | Munglingup, WA | S | 11179 |
| 2003 | Morawa, WA | S | 10937 |
| 2003 | Albany, WA | S | 10981 |
| 2003 | Jerramungup, WA | S | 11185 |
| 2007 | Durham Ox, Vic | S | 07-033 |
| 2009 | Irvingdale, Qld | S | 09-005 |
| 2009 | Mooree, NSW | S | 09-038 |
| 2009 | Muresk, WA | S | M2 |
| 2012 | Boscabel, WA | S | U1 |
| 2012 | Kebaringup, WA | S | U11 |
| 2012 | Lumeah, WA | S | U13 |
| 2012 | Amelup, WA | S | U7, U9 |
| 2013 | Kojonup, WA<sup>B</sup> | S | Ko303, Ko503, Ko603 |
| 2013 | Meckering, WA<sup>B</sup> | S | Meck109, Meck301, Meck305, Meck306, Meck310 |
| 2016 | Gibson, WA | MR1 | 16FRG073 |
| 2017 | Takaralup, WA | HR | 17FRG089 |
| 2017 | Coomalbidgup, WA | MR1 | 17FRG178 |
| 2017 | Takaralup, WA | HR | 17FRG164, 17FRG167 |
| 2017 | Green Range, WA | HR | 17FRG186, 17FRG198 |
| 2018 | Munglinup, WA | MR1 | 18FRG002, 18FRG003, 18FRG005 |
| 2018 | Takaralup, WA | HR | 18FRG007, 18FRG012 |
| 2018 | South Stirling, WA | HR | 18FRG014, 18FRG024 |
| 2018 | Takaralup, WA | HR | 18FRG027, 18FRG033 |
| 2018 | Frankland, WA | HR | 18FRG061 |
| 2018 | Amelup, WA | HR | 18FRG062 |
| 2018 | Dalyup, WA | HR | 18FRG063, 18FRG064, 18FRG065, 18FRG066 |
| 2018 | Gibson, WA | HR | 18FRG180, 18FRG181 |
| 2018 | Dalyup, WA | HR | 18FRG187, 18FRG188, 18FRG189, 18FRG190 |
| 2018 | Pithara, WA | MR2 | 18FRG195 |
| 2019 | Broomehill, WA | HR | 19FRG001, 19FRG002, 19FRG003 |

Location: WA= Western Australia, NSW= New South Wales, Qld= Queensland, Vic= Victoria. Phenotype: S= sensitive to DMIs, MR1= moderately resistant to DMIs (phenotype 1), MR2= moderately resistant to DMIs (phenotype 2), HR= highly resistant to DMIs .<sup>B</sup> Bait trial.

### Discriminatory dose screening

For each isolate, three biological replicates of 4 mm-diameter mycelial plugs taken from the colony edge of 5 day monoconidial cultures grown on V8PDA plates, were inoculated to Yeast Bacto Acetate agar (10 g yeast extract, 10 g Bacto peptone, 10 g sodium acetate, 15 g agar in 1 L H_2_O; YBA), amended with solutions of technical-grade tebuconazole dissolved in ethanol, to a final concentration of 10 µg mL^-1^. An equivalent volume of ethanol was added for the zero fungicide control. Cultures were incubated in darkness at room temperature for 96 h. Isolates were considered to be DMI-sensitive when there was complete cessation of growth on fungicide-amended plates, with isolates considered to be DMI-resistant where the presence of hyphae was observed on surface of plates amended with 10 µg mL^-1^ tebuconazole.

### Characterisation of *in vitro* fungicide sensitivity phenotypes

Fungicide sensitivity of isolates was determined *in vitro* by microtiter assay. Serial dilutions of technical grade fungicides difenoconazole, epoxiconazole, metconazole, prochloraz, propiconazole prothioconazole and tebuconazole, were dissolved in ethanol. For each compound the range of doses tested was adjusted based on the response of the individual isolate. Fungicide stock (or ethanol solvent for the zero fungicide control) was added to YBA medium (10 g yeast extract, 10 g Bacto peptone, 10 g sodium acetate in 1 L deionized H_2_O) at a final concentration 0.5% (v/v); and 95 µL of the resulting mixture dispensed per well to a 96-well tissue culture plate (TrueLine,).

For each isolate, at least two biological replicates of 4 mm-diameter mycelial plugs of monoconidial cultures were inoculated to Barley Leaf Agar (70 g fresh homogenised barley leaves, 20 g agar in 1 L deionised H_2_O; BLA). Sporulation was induced on 5 day cultures as described above. Spores were harvested with sterile deionized H_2_O using a pipette tip and glass spreader, and the resulting suspension filtered through a single layer of sterile cheesecloth and concentrated by centrifugation at 3220 g for 10 min at 4 °C. Spore density was estimated using a haemocytometer and adjusted to 5 × 10^3^ spores mL^−1^; each well of the microtiter was inoculated with 5 μL of suspension to a final concentration of 250 spores mL^−1^. At least three technical replicate wells were inoculated per biological replicate per isolate.

Optical density (OD) was measured at 405 nm with Synergy HT microplate reader (BioTek Instruments, Winooski, VT, USA), using area scan settings. Microtiter plates were measured immediately following inoculation and again at 96 h post-inoculation. EC_50_ values were calculated as previously described (Mair, Deng, et al. 2016). Resistance factors (RF) were calculated as a ratio of the isolate EC_50_ to the mean EC_50_ of isolates collected 1996-2013. EC_50_ values were log_10_-transformed (Liang et al. 2015) prior to statistical analysis with SPSS Statistics 24 (IBM, Armonk, NY, USA), using Kruskal-Wallis H test & Dunnett’s T3.

### DNA extraction and sequencing of *Cyp51A* and *Cyp51B* genes

Mycelia taken from V8PDA plates was snap frozen in liquid nitrogen and homogenised with tungsten carbide beads for 2 min at 30 Hz in a Mixer Mill MM400 (Retsch GmbH, Haan, Germany). DNA was extracted using BioSprint 15 DNA Plant Kit (QIAGEN, Hilden, Germany) and the KingFisher mL Purification System (Thermo Fisher Scientific, Waltham, MA, USA), as per manufacturers’ instructions.

For sequencing the *Cyp51A* and *Cyp51B* promoters, PCR amplification was carried out in 50 μL reaction volume containing 0.25 μL MyTaq DNA Polymerase (5 U μL^−1^; Bioline, London, UK), 10 μL 5 × reaction buffer, 2.5 μL each forward and reverse primer (5 μM; Table 2), 1 μL genomic DNA template (100 ng μL^−1^) and 33.75 μL H_2_O. Thermal cycling was conducted using Mastercycler Nexus (Eppendorf, Hamburg, Germany) with the following parameters: initial denaturation at 95 °C for 2 min, followed by 36 cycles of 95 °C for 30 s, 57.1 °C for 30 s, and 72 °C for 1 min, followed by a final extension at 72 °C for 5 min.

**Table 2.**
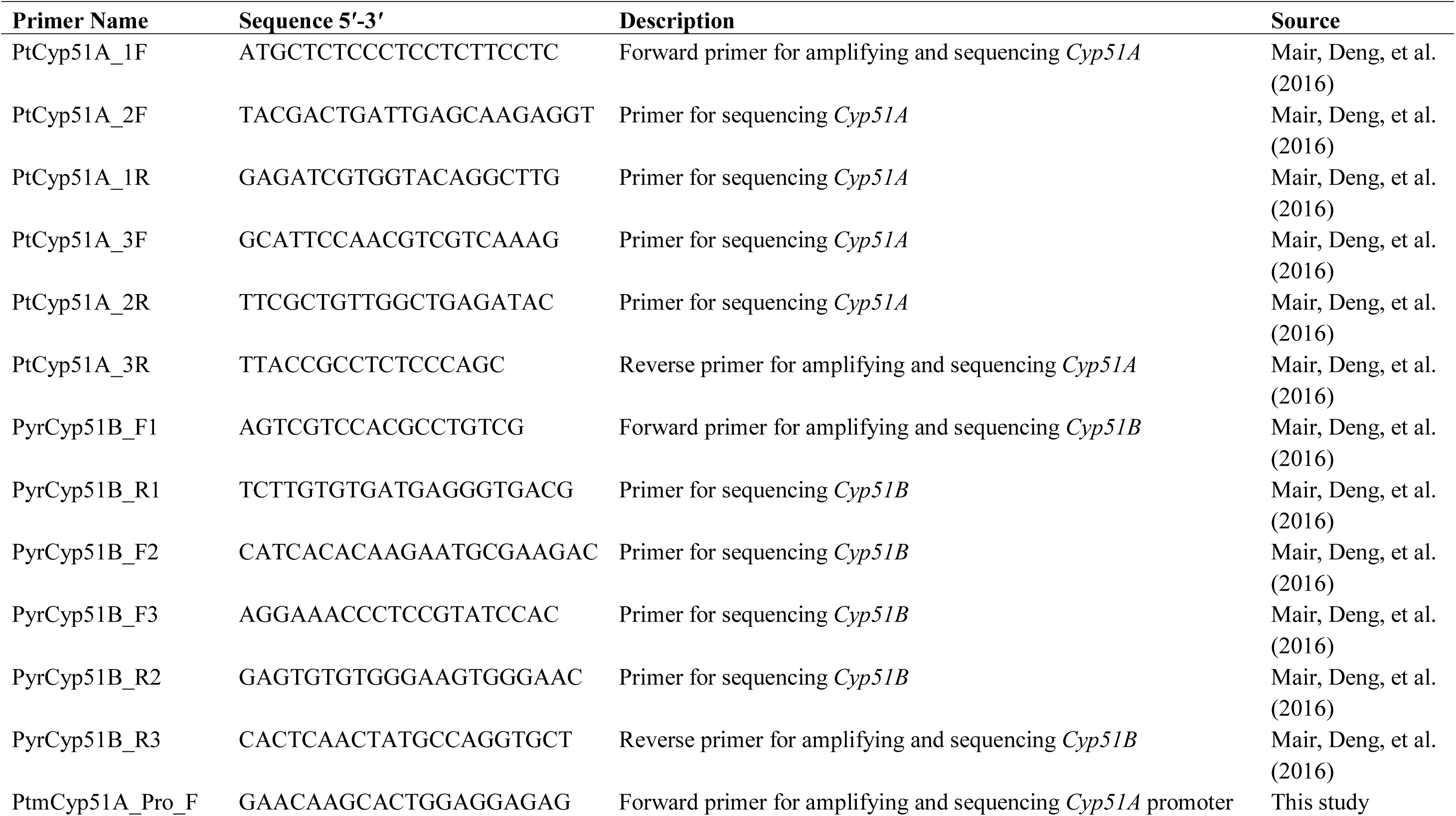
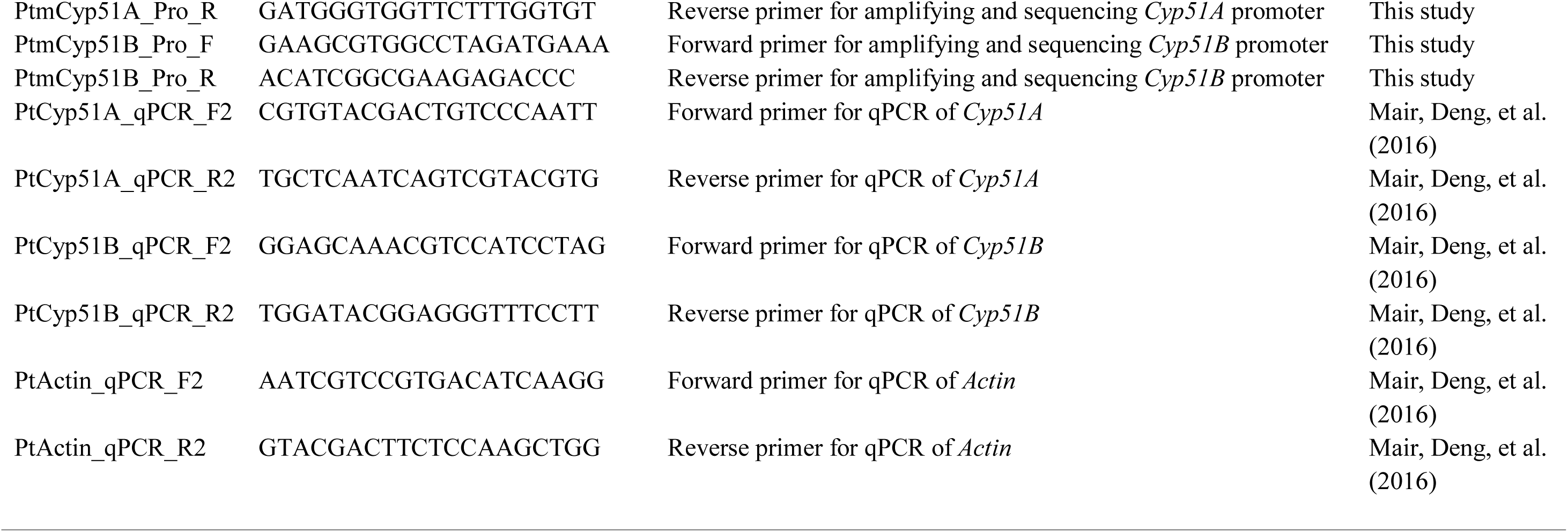
Details of oligonucleotide primers used in this study.

For sequencing the *Cyp51A* coding sequence, PCR amplification was carried out in a 50 μL reaction volume containing 0.5 μL Q5 High Fidelity DNA Polymerase (2 U μL^−1^; New England Biolabs, Ipswich, MA, USA), 10 μL 5 × reaction buffer, 5 μL dNTPs (2 mM), 2.5 μL each forward and reverse primer (10 μM; Table 2), 1 μL genomic DNA template (100 ng μL^−1^) and 28.5 μL H_2_O. Thermal cycling was conducted using Mastercycler Nexus (Eppendorf, Hamburg, Germany) with the following parameters: initial denaturation at 98 °C for 30 s, followed by 36 cycles of 98 °C for 10 s, 70.8 °C for 30 s, and 72 °C for 45 sec, followed by a final extension at 72 °C for 2 min.

For sequencing the *Cyp51B* coding sequence, PCR amplification was carried out in a 50 μL reaction volume containing 0.625 μL DyNAzyme EXT Polymerase (1 U μL^−1^; Thermo Fisher Scientific, Waltham, MA, USA), 5 μL 10 × reaction buffer with 15 mM MgCl_2_, 5 μL dNTPs (2 mM), 2.5 μL each forward and reverse primer (5 μM; Table 2), 0.5 μL genomic DNA template (100 ng μL^−1^) and 33.875 μL H_2_O. Thermal cycling was conducted using Mastercycler Nexus (Eppendorf, Hamburg, Germany) with the following parameters: initial denaturation at 94 °C for 2 min, followed by 38 cycles of 94 °C for 30 s, 60.3 °C for 30 s, and 72 °C for 90 sec, followed by a final extension at 72 °C for 5 min.

PCR products were purified using GenElute PCR Clean-Up Kit (Sigma-Aldrich, St Louis, MO, USA), and sequenced by Macrogen (Seoul, South Korea) using the Applied Biosystems ABI3730XL DNA Analyzer 96-Capillary Array (Thermo Fisher Scientific, Waltham, MA, USA). Sequences were assembled using Geneious 6.1.8 (Biomatters, Auckland, New Zealand) and aligned with ClustalW algorithm (Thompson, Higgins, and Gibson 1994) with the following parameters: IUB scoring matrix, gap opening penalty 15, gap extension penalty 6.66 and free end gaps. Insertion sequences were analysed using BLASTN 2.10.0+ algorithm (Altschul et al. 1990) against *Ptm* isolate SG1 reference genome (GenBank BioProject number |PRJEB18107)(Syme et al. 2018). Prediction of promoter sequences within the upstream regulatory regions performed using Neural Network Promoter Prediction 2.2 software (http://www.fruitfly.org/seq_tools/promoter.html) with a cutoff score of 0.8 (Reese 2001). Prediction of fungal transcription factor binding sites within insertion sequences performed with PROMO 3.0.2 (http://alggen.lsi.upc.es/cgi-bin/promo_v3/promo/promoinit.cgi?dirDB=TF_8.3).

### Growth of fungal cultures and RNA extraction for gene expression analysis

To determine the growth rate of fungal cultures, six biological replicates of a 4 mm-diameter mycelial plug taken from the colony edge of a 5 day culture grown on V8PDA were inoculated to 5 mL of Fries2 media (Fries 1938) in a 6-well cell culture plate (Corning, NY, USA) and incubated at 22 °C, 150 rpm in darkness. OD at 405 nm was measured in hourly intervals with a Synergy HT microplate reader (BioTek Instruments, Winooski, VT, USA) using area scan settings. At 76 h post-inoculation, when cultures were in the exponential phase of growth (data not shown), solutions of tebuconazole, epoxiconazole or prochloraz dissolved in ethanol were added to the culture media at a final concentration equivalent to the respective EC_50_ of each isolate as described by Cools et al. (2012); an equivalent volume of ethanol solvent was added for the zero fungicide controls. At 100 h post-inoculation, cultures were harvested by centrifugation, snap frozen in liquid nitrogen and freeze-dried. Tissue was homogenised with tungsten carbide beads as described above, and RNA extracted using Spectrum Plant Total RNA Kit (Sigma-Aldrich, St Louis, MO, USA). RNA extracts were treated with Invitrogen TURBO DNA-free Kit (Thermo Fisher Scientific, Waltham, MA, USA) and cDNA synthesised using iScript cDNA Synthesis Kit (Bio-Rad Laboratories, Hercules, CA, USA).

### Quantitative PCR for gene expression analysis

Quantitative PCR was conducted on CFX96 C1000 Real-Time System (Bio-Rad Laboratories, Hercules, CA, USA), using iTaq Universal SYBR Green SuperMix (Bio-Rad Laboratories, Hercules, CA, USA) as per manufacturer’s instructions and primers listed in Table 2. PCR parameters were initial denaturation at 95 °C for 5 min, followed by 40 cycles of 95 °C for 10 s, 58 °C for 15 s, and 72 °C for 15 s, followed by a melt curve at 0.5 °C increments of 5 s each between 65 °C and 95 °C.

Mean fold change of gene expression was calculated by 2^−ΔΔCt^ (Livak and Schmittgen 2001) relative to constitutive expression in isolate M2 (ethanol-treated control), normalized to *Actin* (NCBI accession no. XM_003298028) as the endogenous control gene, with at least 3 biological replicates and four technical replicates per biological replicate. Statistical analysis of 2^−ΔΔCt^ data was performed with SPSS Statistics 24 (IBM, Armonk, NY, USA) using Kruskal-Wallis H test & Dunnett’s T3.

## RESULTS

### *In vitro* characterisation reveals different DMI fungicide sensitivity phenotypes in *Ptm*

Twenty-two monoconidial *Ptm* isolates collected between 1996 and 2013 were tested by microtiter assay to determine 50% effective concentrations (EC_50_) to the DMI fungicides tebuconazole, epoxiconazole and prothioconazole (Table 3). Isolate EC_50_ values for tebuconazole ranged from 0.04 to 0.69 µg mL^-1^ (mean 0.34 µg mL^-1^); for epoxiconazole 0.02 to 0.37 µg mL^-1^ (mean 0.18 µg mL^-1^); and for prothioconazole 0.03 to 0.14 µg mL^-1^ (mean 0.07 µg mL^-1^). A subset of three sensitive isolates (SG1, M2 and U7), were also characterised for EC_50_ values to the DMI compounds propiconazole (0.09-0.24 µg mL^-1^, mean 0.19 µg mL^-1^), metconazole (0.12-0.39 µg mL^-1^, mean 0.29 µg mL^-1^), difenoconazole (0.011-0.044 µg mL^-1^, mean 0.029 µg mL^-1^), and prochloraz (0.011-0.038 µg mL^-1^, mean 0.027 µg mL^-1^) (Table 4).

**Table 3.** EC_50_ of *Pyrenophora teres* f. sp. *maculata* isolates to tebuconazole, epoxiconazole and prothioconazole.

| Isolate | Tebuconazole |  | Epoxiconazole |  | Prothioconazole |  |
| --- | --- | --- | --- | --- | --- | --- |
|  | EC<sub>50</sub> (µg mL<sup>-1</sup>) | RF<sup>a</sup> | EC<sub>50</sub> (µg mL<sup>-1</sup>) | RF | EC<sub>50</sub> (µg mL<sup>-1</sup>) | RF |
| **07-033** | 0.35 (±0.08)<sup>b</sup> |  | 0.16 (±0.01) |  | 0.067 (±0.007) |  |
| **09-005** | 0.32 (±0.07) |  | 0.14 (±0.02) |  | 0.041 (±0.003) |  |
| **09-038** | 0.13 (±0.03) |  | 0.06 (±0.01) |  | 0.041 (±0.010) |  |
| **10937** | 0.11 (±0.05) |  | 0.04 (±0.01) |  | 0.045 (±0.014) |  |
| **10981** | 0.48 (±0.08) |  | 0.29 (±0.03) |  | 0.081 (±0.001) |  |
| **11179** | 0.43 (±0.03) |  | 0.20 (±0.02) |  | 0.079 (±0.012) |  |
| **11185** | 0.39 (±0.06) |  | 0.15 (±0.02) |  | 0.080 (±0.008) |  |
| **Ko303** | 0.53 (±0.10) |  | 0.26 (±0.05) |  | 0.068 (±0.003) |  |
| **Ko503** | 0.16 (±0.05) |  | 0.29 (±0.09) |  | 0.053 (±0.025) |  |
| **Ko603** | 0.33 (±0.03) |  | 0.10 (±0.02) |  | 0.110 (±0.030) |  |
| **Meck109** | 0.44 (±0.08) |  | 0.16 (±0.04) |  | 0.093 (±0.016) |  |
| **Meck301** | 0.50 (±0.17) |  | 0.12 (±0.04) |  | 0.091 (±0.022) |  |
| **Meck305** | 0.17 (±0.02) |  | 0.14 (±0.02) |  | 0.045 (±0.001) |  |
| **Meck306** | 0.19 (±0.03) |  | 0.19 (±0.04) |  | 0.082 (±0.002) |  |
| **Meck310** | 0.24 (±0.03) |  | 0.20 (±0.06) |  | 0.058 (±0.023) |  |
| **U1** | 0.33 (±0.05) |  | 0.21 (±0.05) |  | 0.084 (±0.033) |  |
| **U11** | 0.34 (±0.04) |  | 0.11 (±0.03) |  | 0.105 (±0.024) |  |
| **U13** | 0.41 (±0.03) |  | 0.08 (±0.02) |  | 0.028 (±0.001) |  |
| **U9** | 0.04 (±0.01) |  | 0.02 (±0.008) |  | 0.053 (±0.019) |  |
| **U7** | 0.48 (±0.06) |  | 0.31 (±0.03) |  | 0.104 (±0.027) |  |
| **M2** | 0.69 (±0.04) |  | 0.37 (±0.03) |  | 0.142 (±0.011) |  |
| **SG1** | 0.37 (±0.05) |  | 0.26 (±0.03) |  | 0.078 (±0.009) |  |
|  | **M= 0.34** |  | **M= 0.18** |  | **M= 0.074** |  |
| **16FRG073** | 2.10 (±0.15) | 6.2 | 1.31 (±0.17) | 7.4 | 0.35 (±0.06) | 4.7 |
| **17FRG178** | 2.16 (±0.30) | 6.4 | 2.18 (±0.17) | 12.4 | 0.73 (±0.10) | 9.8 |
| **18FRG002** | 3.57 (±0.37) | 10.6 | 2.32 (±0.18) | 13.2 | 0.95 (±0.15) | 12.9 |
| **18FRG003** | 2.53 (±0.45) | 7.5 | 1.41 (±0.19) | 8.0 | 0.29 (±0.03) | 3.9 |
| **18FRG005** | 1.98 (±0.28) | 5.9 | 1.40 (±0.22) | 7.9 | 0.67 (±0.10) | 9.1 |

|  | ***M* = 2.47** | ***M* = 7.3** | ***M* = 1.72** | ***M* = 9.8** | ***M* = 0.60** | ***M* = 8.1** |
| --- | --- | --- | --- | --- | --- | --- |
| **18FRG195** | 2.04 (±0.24) | 6.0 | 0.09 (±0.02) | 0.5 | 0.20 (±0.03) | 2.7 |
| **17FRG089** | 15.0 (±1.0) | 44.4 | 1.35 (±0.21) | 7.6 | 2.02 (±0.23) | 27.3 |
| **17FRG164** | 17.7 (±0.7) | 52.3 | 2.14 (±0.29) | 12.1 | 1.66 (±0.13) | 22.4 |
| **17FRG167** | 16.1 (±1.8) | 47.5 | 1.88 (±0.26) | 10.7 | 2.28 (±0.30) | 30.8 |
| **17FRG186** | 18.4 (±0.9) | 54.4 | 1.76 (±0.39) | 10.0 | 2.14 (±0.53) | 29.0 |
| **17FRG198** | 19.4 (±1.1) | 57.4 | 2.36 (±0.29) | 13.4 | 2.70 (±0.32) | 36.5 |
| **18FRG007** | 17.9 (±1.2) | 52.9 | 2.27 (±0.30) | 12.9 | 3.28 (±0.45) | 44.4 |
| **18FRG012** | 16.5 (±1.5) | 48.8 | 2.15 (±0.41) | 12.2 | 1.69 (±0.16) | 22.9 |
| **18FRG014** | 18.5 (±1.8) | 54.9 | 1.29 (±0.23) | 7.3 | 1.31 (±0.26) | 17.8 |
| **18FRG024** | 19.4 (±1.1) | 57.5 | 1.80 (±0.28) | 10.2 | 2.49 (±0.41) | 33.7 |
| **18FRG027** | 17.8 (±1.9) | 52.6 | 1.87 (±0.30) | 10.6 | 1.17 (±0.38) | 15.8 |
| **18FRG033** | 19.0 (±1.7) | 56.3 | 1.67 (±0.38) | 9.4 | 0.99 (±0.21) | 13.3 |
| **18FRG063** | 10.3 (±1.7) | 30.5 | 0.84 (±0.23) | 4.8 | 0.75 (±0.22) | 10.1 |
| **18FRG064** | 15.7 (±0.9) | 46.3 | 1.00 (±0.11) | 5.7 | 1.10 (±0.25) | 14.9 |
| **18FRG065** | 14.7 (±0.8) | 43.6 | 1.20 (±0.12) | 6.8 | 1.21 (±0.25) | 16.3 |
| **18FRG066** | 14.0 (±2.2) | 41.4 | 1.73 (±0.20) | 9.8 | 1.13 (±0.24) | 15.2 |
| **19FRG001** | 18.3 (±1.6) | 54.2 | 0.80 (±0.14) | 4.5 | 1.54 (±0.30) | 20.8 |
| **19FRG002** | 21.8 (±1.5) | 64.5 | 1.42 (±0.21) | 8.1 | 2.60 (±0.36) | 35.2 |
|  | ***M* = 17.1** | ***M* = 50.6** | ***M* = 1.62** | ***M* = 9.2** | ***M* = 1.77** | ***M* = 23.9** |

EC<sub>50</sub> values are the mean of at least two biological replicates. <sup>a</sup>Resistance Factor; <sup>b</sup>(± Standard error of the mean).

**Table 4.**
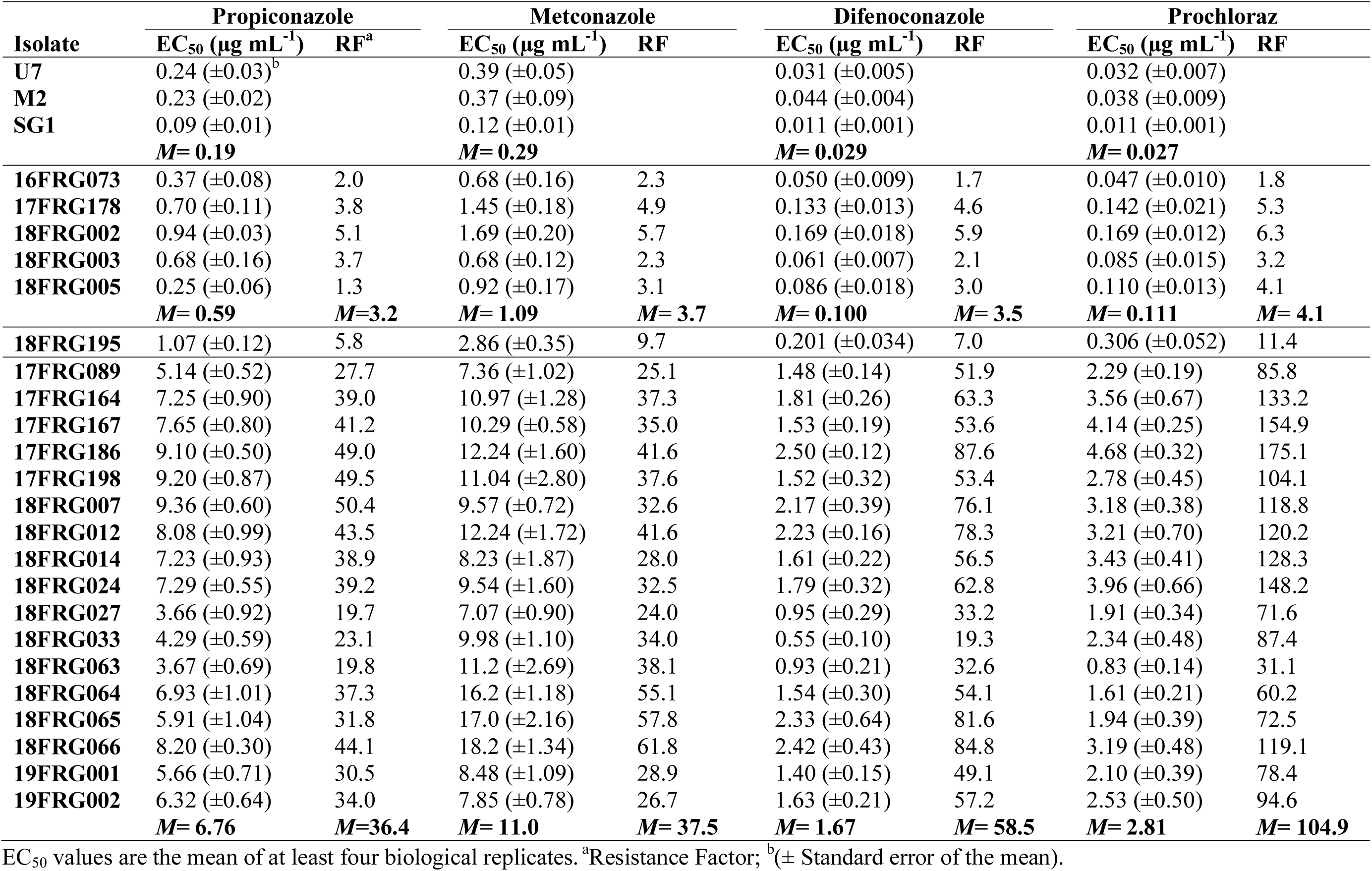
EC_50_ of *Pyrenophora teres* f.sp. *maculata* isolates to propiconazole, metconazole, difenoconazole and prochloraz.

| Isolate | Propiconazole |  | Metconazole |  | Difenoconazole |  | Prochloraz |  |
| --- | --- | --- | --- | --- | --- | --- | --- | --- |
|  | EC<sub>50</sub> (µg mL<sup>-1</sup>) | RF<sup>a</sup> | EC<sub>50</sub> (µg mL<sup>-1</sup>) | RF | EC<sub>50</sub> (µg mL<sup>-1</sup>) | RF | EC<sub>50</sub> (µg mL<sup>-1</sup>) | RF |
| U7 | 0.24 (±0.03)<sup>b</sup> |  | 0.39 (±0.05) |  | 0.031 (±0.005) |  | 0.032 (±0.007) |  |
| M2 | 0.23 (±0.02) |  | 0.37 (±0.09) |  | 0.044 (±0.004) |  | 0.038 (±0.009) |  |
| SG1 | 0.09 (±0.01) |  | 0.12 (±0.01) |  | 0.011 (±0.001) |  | 0.011 (±0.001) |  |
|  | **M= 0.19** |  | **M= 0.29** |  | **M= 0.029** |  | **M= 0.027** |  |
| 16FRG073 | 0.37 (±0.08) | 2.0 | 0.68 (±0.16) | 2.3 | 0.050 (±0.009) | 1.7 | 0.047 (±0.010) | 1.8 |
| 17FRG178 | 0.70 (±0.11) | 3.8 | 1.45 (±0.18) | 4.9 | 0.133 (±0.013) | 4.6 | 0.142 (±0.021) | 5.3 |
| 18FRG002 | 0.94 (±0.03) | 5.1 | 1.69 (±0.20) | 5.7 | 0.169 (±0.018) | 5.9 | 0.169 (±0.012) | 6.3 |
| 18FRG003 | 0.68 (±0.16) | 3.7 | 0.68 (±0.12) | 2.3 | 0.061 (±0.007) | 2.1 | 0.085 (±0.015) | 3.2 |
| 18FRG005 | 0.25 (±0.06) | 1.3 | 0.92 (±0.17) | 3.1 | 0.086 (±0.018) | 3.0 | 0.110 (±0.013) | 4.1 |
|  | **M= 0.59** | **M=3.2** | **M= 1.09** | **M= 3.7** | **M= 0.100** | **M= 3.5** | **M= 0.111** | **M= 4.1** |
| 18FRG195 | 1.07 (±0.12) | 5.8 | 2.86 (±0.35) | 9.7 | 0.201 (±0.034) | 7.0 | 0.306 (±0.052) | 11.4 |
| 17FRG089 | 5.14 (±0.52) | 27.7 | 7.36 (±1.02) | 25.1 | 1.48 (±0.14) | 51.9 | 2.29 (±0.19) | 85.8 |
| 17FRG164 | 7.25 (±0.90) | 39.0 | 10.97 (±1.28) | 37.3 | 1.81 (±0.26) | 63.3 | 3.56 (±0.67) | 133.2 |
| 17FRG167 | 7.65 (±0.80) | 41.2 | 10.29 (±0.58) | 35.0 | 1.53 (±0.19) | 53.6 | 4.14 (±0.25) | 154.9 |
| 17FRG186 | 9.10 (±0.50) | 49.0 | 12.24 (±1.60) | 41.6 | 2.50 (±0.12) | 87.6 | 4.68 (±0.32) | 175.1 |
| 17FRG198 | 9.20 (±0.87) | 49.5 | 11.04 (±2.80) | 37.6 | 1.52 (±0.32) | 53.4 | 2.78 (±0.45) | 104.1 |
| 18FRG007 | 9.36 (±0.60) | 50.4 | 9.57 (±0.72) | 32.6 | 2.17 (±0.39) | 76.1 | 3.18 (±0.38) | 118.8 |
| 18FRG012 | 8.08 (±0.99) | 43.5 | 12.24 (±1.72) | 41.6 | 2.23 (±0.16) | 78.3 | 3.21 (±0.70) | 120.2 |
| 18FRG014 | 7.23 (±0.93) | 38.9 | 8.23 (±1.87) | 28.0 | 1.61 (±0.22) | 56.5 | 3.43 (±0.41) | 128.3 |
| 18FRG024 | 7.29 (±0.55) | 39.2 | 9.54 (±1.60) | 32.5 | 1.79 (±0.32) | 62.8 | 3.96 (±0.66) | 148.2 |
| 18FRG027 | 3.66 (±0.92) | 19.7 | 7.07 (±0.90) | 24.0 | 0.95 (±0.29) | 33.2 | 1.91 (±0.34) | 71.6 |
| 18FRG033 | 4.29 (±0.59) | 23.1 | 9.98 (±1.10) | 34.0 | 0.55 (±0.10) | 19.3 | 2.34 (±0.48) | 87.4 |
| 18FRG063 | 3.67 (±0.69) | 19.8 | 11.2 (±2.69) | 38.1 | 0.93 (±0.21) | 32.6 | 0.83 (±0.14) | 31.1 |
| 18FRG064 | 6.93 (±1.01) | 37.3 | 16.2 (±1.18) | 55.1 | 1.54 (±0.30) | 54.1 | 1.61 (±0.21) | 60.2 |
| 18FRG065 | 5.91 (±1.04) | 31.8 | 17.0 (±2.16) | 57.8 | 2.33 (±0.64) | 81.6 | 1.94 (±0.39) | 72.5 |
| 18FRG066 | 8.20 (±0.30) | 44.1 | 18.2 (±1.34) | 61.8 | 2.42 (±0.43) | 84.8 | 3.19 (±0.48) | 119.1 |
| 19FRG001 | 5.66 (±0.71) | 30.5 | 8.48 (±1.09) | 28.9 | 1.40 (±0.15) | 49.1 | 2.10 (±0.39) | 78.4 |
| 19FRG002 | 6.32 (±0.64) | 34.0 | 7.85 (±0.78) | 26.7 | 1.63 (±0.21) | 57.2 | 2.53 (±0.50) | 94.6 |
|  | **M= 6.76** | **M=36.4** | **M= 11.0** | **M= 37.5** | **M= 1.67** | **M= 58.5** | **M= 2.81** | **M= 104.9** |

EC<sub>50</sub> values are the mean of at least four biological replicates. <sup>a</sup>Resistance Factor; <sup>b</sup>(± Standard error of the mean).

As outlined previously (Mair, Deng, et al. 2016), and based on the above results, a cut-off concentration of 10 µg mL^-1^ tebuconazole was determined to completely inhibit the growth of putatively DMI-sensitive isolates. Using this threshold an additional 266 isolates collected between 2014-2019 (Table S1) were pre-screened, and from the 2016 season onwards fifty-nine isolates were identified as able to grow on a discriminatory dose of tebuconazole. A subset of twenty-three of these putatively resistant isolates were further analysed by microtiter test for sensitivity to the seven DMI compounds listed above (Tables 3 and 4).

EC_50_ values to the seven DMI compounds assorted strains into several different phenotypic groups. In addition to the isolates from 1998-2013, classified as sensitive (S), the isolates able to grow on a discriminatory dose of tebuconazole were classified as moderately-resistant (two phenotypic groups identified: MR1 and MR2) or highly-resistant (HR) to DMIs. Five strains (16FRG073, 17FRG178, 18FRG002, 18FRG003, and 18FRG005) were classified as MR1, having EC_50_ values higher than S isolates for the compounds difenoconazole (0.061-0.169 µg mL^-1^, mean 0.100 µg mL^-1^), epoxiconazole (1.31-2.32 µg mL^-1^, mean 1.72 µg mL^-1^), metconazole (0.68-1.69 µg mL^-1^, mean 1.09 µg mL^-1^), prochloraz (0.047-0.169 µg mL^-1^, mean 0.111 µg mL^-1^), propiconazole (0.25-0.94 µg mL^-1^, mean 0.59 µg mL^-1^), prothioconazole (0.29-0.95 µg mL^-1^, mean 0.60 µg mL^-1^), and tebuconazole (1.98-3.57 µg mL^-1^, mean 2.47 µg mL^-1^). One strain (18FRG195), classified as MR2, had EC_50_ values higher than S isolates for the compounds difenoconazole (0.201 µg mL^-1^), metconazole (2.86 µg mL^-1^), prochloraz (0.306 µg mL^-1^), propiconazole (1.07 µg mL^-1^), prothioconazole (0.20 µg mL^-1^), and tebuconazole (2.04 µg mL^-1^), but lower than S isolates for epoxiconazole (0.09 µg mL^-1^). The remaining 17 strains were classified as having an HR phenotype, EC_50_ values were higher than S, MR1, and MR2 for the compounds difenoconazole (0.55-2.50 µg mL^-1^, mean 1.67 µg mL^-1^), metconazole (7.07-18.2 µg mL^-1^, mean 11.0 µg mL^-1^), prochloraz (0.83-4.68 µg mL^-1^, mean 2.81 µg mL^-1^), propiconazole (3.66-9.36 µg mL^-1^, mean 6.76 µg mL^-1^), prothioconazole (0.75-3.28 µg mL^-1^, mean 1.77 µg mL^-1^), and tebuconazole (10.3-21.8 µg mL^-1^, mean 17.1 µg mL^-1^), and higher than that of S and MR2, but similar to MR1, for epoxiconazole (0.80-2.36 µg mL^-1^, mean 1.62 µg mL^-1^).

A Kruskal-Wallis test showed significant differences at *P* < .05 among the four phenotypic groups for difenoconazole (*H*(3) = 145.874, *P* < .001), epoxiconazole (*H*(3) = 194.240, *P* < .001), metconazole (*H*(3) = 151.042, *P* < .001), prochloraz (*H*(3) = 144.640, *P* < .001), propiconazole (*H*(3) = 143.016, *P* < .001), prothioconazole (*H*(3) = 199.680, *P* < .001) and tebuconazole (*H*(3) = 228.617, *P* < .001). Post hoc comparisons with Dunnett’s T3 test showed significant (*P* < .001) differences among each of the groups for tebuconazole, except for between MR1 and MR2 where the difference was not significant (*P* = .766). For epoxiconazole, Dunnett’s T3 showed significant (*P* < .05) differences among each of the groups, except for between MR1 and HR which did not differ significantly (*P* = .358). Post hoc comparisons also showed significant differences among each of the four groups for difenoconazole (*P* < .02), metconazole (*P* < .001), prochloraz (*P* < .001), propiconazole (*P* < .002), and prothioconazole (*P* < .002).

### DMI fungicide resistance in *Ptm* is correlated with presence of several different point mutations resulting in F489L (*F495L*) in CYP51A

Of the paralogous genes encoding the DMI target, *Cyp51B* (1.68 kbp) was sequenced in five sensitive isolates (11185, 09-005, 09-038, M2 and U7) and the fifty-nine isolates identified as having reduced sensitivity to DMI fungicides. A single allele of *Cyp51B* was detected across all the isolates tested, identical to that of the reference sequence from isolate SG1 (GenBank accession JQ314399). Sequencing of the *Cyp51A* paralog (1.512 kbp) in three sensitive isolates (SG1, M2 and U7) and the fifty-nine DMI-reduced sensitive isolates revealed five distinct alleles (Table 5, Figure S1). Sensitive isolates SG1 and U7, and MR1 isolates 16FRG073, 17FRG178, 18FRG002, 18FRG003 and 18FRG005, share the allele *Sg1-A* (GenBank accession …). Sensitive isolate M2 carries the allele *M2-A* (GenBank accession …), which differs from *Sg1-A* by nine nucleotides including three non-synonymous changes, resulting in the substitutions C15F, I16T and M104I. Across fifty-three HR isolates tested two alleles were detected. The majority (fifty isolates) shared the allele *17089-A* (GenBank accession …) which differed from *M2-A* by five nucleotides including one non-synonymous mutation. The predicted amino acid sequence of *17089-A* was identical to *M2-A* except for the substitution of phenylalanine by leucine at position 489, resulting from a C to A transversion at position 1467 of the nucleotide sequence (c1467a) (Figure 1, Figure S3). This same c1467a mutation, resulting in the substitution F489L (homologous to the archetypal *F495I* mutation in the CYP51A of *Aspergillus fumigatus*) (Mair, Lopez-Ruiz, et al. 2016), has been previously reported in the closely related pathogen *Pyrenophora teres* f. sp. *teres* (*Ptt*) where it has been correlated with reduced sensitivity to a range of DMI fungicides (Mair, Deng, et al. 2016). The same F489L (*F495L*) amino acid substitution was also observed in the remaining three HR isolates (19FRG001, 19FRG002, 19FRG003), although in this case resulting from a C to G transversion at position 1467 (c1467g). This *19001* allele (GenBank accession …) differed from the wild-types *M2-A* and *Sg1-A* by six and eleven nucleotides respectively, and from *17089-A* by six synonymous changes; the predicted amino acid sequence of *19001-A* was identical to that of *17089-A*. The MR2 isolate 18FRG195 carries a unique allele (*18195-A*), the same F489L (*F495L*) amino acid substitution was again observed in *18195-A*, but in this case resulting from a T to C transition at position 1465 of the nucleotide sequence (t1465c). The *18195-A* allele also carries ten synonymous mutations and three additional non-synonymous mutations compared to the reference *Sg1-A*, resulting in the substitutions L219F, P225L and I253M in the predicted amino acid sequence.

**Figure 1.**
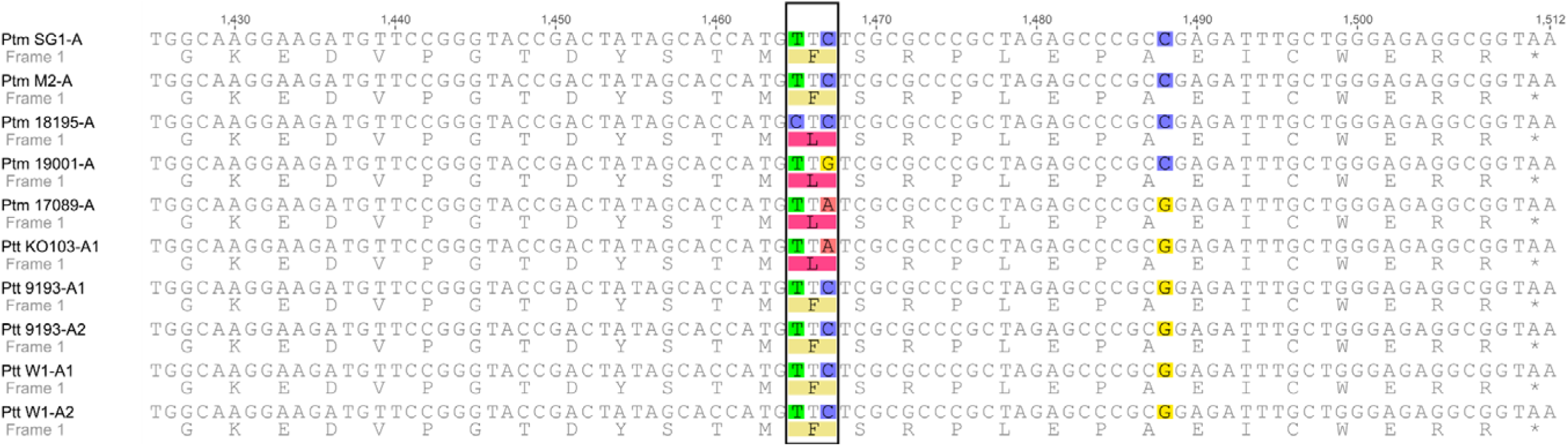
Partial alignment of *Cyp51A* alleles in *Pyrenophora teres* species. 3’ terminal regions of the coding sequences of the five *P. teres* f. sp. *maculata Cyp51A* alleles (*Sg1-A, M2-A, 18195-A, 19001-A, 17089-A*) detected in this study and five *P. teres* f. sp. *teres Cyp51A* alleles (*KO103-A1, 9193-A1, 9193-A2, W1-A1, W1-A2*) reported by Mair, Deng, et al. (2016). Allele *17089-A,* found in highly DMI-resistant (HR) strains, carries a single nucleotide polymorphism c1467a, resulting in the amino acid substitution F489L, identical to the mutation found in resistant *Ptt* (allele *KO103-A1*). Alleles *18195-A,* found in some moderately DMI-resistant strains (MR2), and *19001-A*, found in DMI-HR strains, also carry the F489L substitution, but resultant from the nucleotide polymorphisms t1465c and c1467g, respectively. Position numbers shown above alignment are relative to start codon in sequence of sensitive reference isolate SG1, translated amino acid sequences are shown below nucleotide sequence, polymorphisms in alignment are highlighted, codon 489 boxed. Alignment generated in Geneious version 6.1.8 software (Biomatters) using ClustalW algorithm with IUB scoring matrix, gap opening penalty 15, gap extension penalty 6.66 and free end gaps.

**Table 5.** *Cyp51A* alleles found in *Pyrenophora teres* f. sp. *maculata*.

| Allele | Polymorphisms in amino acid sequence |  |  |  |  |  |  | Found in isolate |
| --- | --- | --- | --- | --- | --- | --- | --- | --- |
|  | 15 | 16 | 104 | 219 | 225 | 253 | 489 |  |
| *Sg1-A* | C | I | M | L | P | I | F | SG1<sup>a</sup>, U7<sup>a</sup>, 16FRG073<sup>b</sup>, 17FRG178<sup>b</sup>, 18FRG002<sup>b</sup>, 18FRG003<sup>b</sup>, 18FRG005<sup>b</sup> |
| *18195-A* | C | I | M | F | L | M | L | 18FRG195<sup>c</sup> |
| *M2-A* | F | T | I | L | P | I | F | M2<sup>a</sup> |
| *17089-A* | F | T | I | L | P | I | L | 17FRG089<sup>d</sup>, 17FRG164<sup>d</sup>, 17FRG165<sup>d</sup>, 17FRG166<sup>d</sup>, 17FRG167<sup>d</sup>, 17FRG186<sup>d</sup>, 17FRG187<sup>d</sup>, 17FRG195<sup>d</sup>, 17FRG196<sup>d</sup>, 17FRG197<sup>d</sup>, 17FRG198<sup>d</sup>, 18FRG007<sup>d</sup>, 18FRG008<sup>d</sup>, 18FRG009<sup>d</sup>, 18FRG010<sup>d</sup>, 18FRG011<sup>d</sup>, 18FRG012<sup>d</sup>, 18FRG013<sup>d</sup>, 18FRG014<sup>d</sup>, 18FRG015<sup>d</sup>, 18FRG016<sup>d</sup>, 18FRG017<sup>d</sup>, 18FRG018<sup>d</sup>, 18FRG019<sup>d</sup>, 18FRG020<sup>d</sup>, 18FRG021<sup>d</sup>, 18FRG022<sup>d</sup>, 18FRG023<sup>d</sup>, 18FRG024<sup>d</sup>, 18FRG025<sup>d</sup>, 18FRG026<sup>d</sup>, 18FRG027<sup>d</sup>, 18FRG028<sup>d</sup>, 18FRG029<sup>d</sup>, 18FRG030<sup>d</sup>, 18FRG031<sup>d</sup>, 18FRG032<sup>d</sup>, 18FRG033<sup>d</sup>, 18FRG061<sup>d</sup>, 18FRG062<sup>d</sup>, 18FRG063<sup>d</sup>, 18FRG064<sup>d</sup>, 18FRG065<sup>d</sup>, 18FRG066<sup>d</sup>, 18FRG180<sup>d</sup>, 18FRG181<sup>d</sup>, 18FRG187<sup>d</sup>, 18FRG188<sup>d</sup>, 18FRG189<sup>d</sup>, 18FRG190<sup>d</sup> |
| *19001-A<sup>e</sup>* | F | T | I | L | P | I | L | 19FRG001<sup>d</sup>, 19FRG002<sup>d</sup>, 19FRG003<sup>d</sup> |

<sup>a</sup>DMI-S, <sup>b</sup>DMI-MR1, <sup>c</sup>DMI-MR2, <sup>d</sup>DMI-HR, <sup>e</sup>Predicted amino acid sequence identical to *17089-A*, coding sequence differs from *17089-A* by six synonymous nucleotide changes.

### DMI fungicide resistance in *Ptm* is correlated with the presence of insertion elements at several different positions in the *Cyp51A* upstream region

The region extending 463 bp upstream of the start codon of *Cyp51B* was sequenced in seven sensitive isolates (SG1, 10981, 11185, 09-005, 09-038, M2 and U7) and the fifty-nine isolates identified as having reduced sensitivity to DMI fungicides. No changes in the upstream region of *Cyp51B* were observed among any of the isolates. The region extending 898 bp upstream of the start codon of *Cyp51A* was sequenced in three sensitive isolates (SG1, M2 and U7) and the fifty-nine isolates identified as having reduced sensitivity to DMI fungicides, seven different alleles were detected (Figure 2, Figure S2). The sensitive isolates SG1 and U7, and the MR2 isolate 18FRG195 shared an identical upstream sequence (*Sg1-Ap*, GenBank accession …). The sequence from the sensitive isolate M2 differed from the SG1 upstream region by eight single nucleotide polymorphisms at –460, –360, –261, –117, –72, – 71, –70, and –64 (*M2*-*Ap*, GenBank accession …). A 134 bp insertion at –74 to the start codon (labelled *PtTi-1*) was observed in the MR1 isolates 16FRG073 and 18FRG002, the sequence was otherwise identical to that of SG1 (Genbank accession …). In the MR1 isolates 17FRG178 and 18FRG005, a 134 bp insertion was found at –75 to the start codon (*PtTi-3*), the sequence was otherwise identical to that of SG1 (Genbank accession …). In MR1 isolate 18FRG003, a 134 bp insertion was located at –66 to the start codon (*PtTi-4*), the sequence was otherwise identical to that of SG1 (Genbank accession …). In fifty HR isolates carrying the c1467a (F489L) substitution (Figure S3), a 134 bp insertion element was found at –46 to the start codon (*PtTi-2*), the sequence was otherwise identical to that of M2 (Genbank accession …). In three HR isolates carrying the c1467g (F489L) substitution, a 134 bp insertion element was found at –90 to the start codon (*PtTi-5*), the sequence was otherwise identical to that of M2 (Genbank accession …).

**Figure 2.**
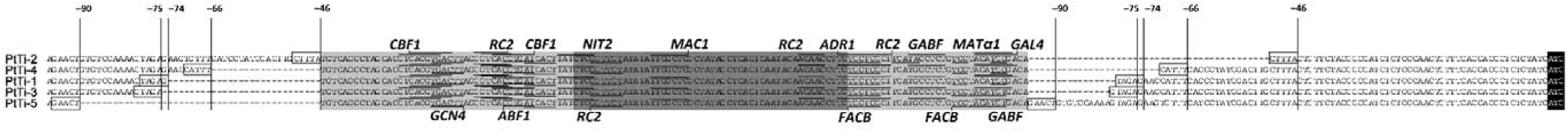
Alignment of upstream region of *PtmCyp51A* in DMI-MR1 & DMI-HR isolates. Regions upstream of *Cyp51A* in highly DMI-resistant (HR) and some moderately DMI-resistant (MR1) strains carry 134 bp insertions relative to sequence from sensitive strain SG1. The insertion were found at –46 (*PtTi-2*), –66 (*PtTi-4*), –74 (*PtTi-1*), –75 (*PtTi-3*), or – 90 (*PtTi-5*) to the start codon of *Cyp51A*. The insertion elements share >95% sequence identity, with a 129 bp consensus sequence homologous to the Long Terminal Repeats (LTRs) of *Ty1*/*Copia* LTR retrotransposons found in the reference genome of SG1. The consensus sequence is flanked by 5 bp direct repeats, presumed target site duplications (TSDs). The insertion elements all contained an identical predicted promoter sequence with a score of 0.99, and a number of predicted transcription factor binding sites. Light grey box: sequences homologous to *Ty1*/*Copia* LTR retrotransposons; Dark grey box: predicted promoter sequences; White box: putative TSD sequences; Black box: start codons; Underlined/overlined: predicted transcription factor binding sites. Position numbers shown above alignment are relative to start codon of *Cyp51A* in sequence of sensitive reference isolate SG1 (not shown in alignment). Promoter prediction performed with Neural Network Promoter Prediction 2.2 software (http://www.fruitfly.org/seq_tools/promoter.html, Reese 2001). Transcription factor binding sites predicted with PROMO 3.0.2 (http://alggen.lsi.upc.es/cgi-bin/promo_v3/promo/promoinit.cgi?dirDB=TF_8.3). Alignment generated in Geneious version 6.1.8 software (Biomatters) using ClustalW algorithm with IUB scoring matrix, gap opening penalty 15, gap extension penalty 6.66 and free end gaps.

Sequence analysis showed that the five 134 bp insertion elements shared at least 95.5% identity. Blastn analysis against the SG1 genome revealed that for each of the five insertion elements, 56 similar sequences were found in the sensitive reference isolate, with E values ranging from 1.13e-63 to 1.13e-06 and pairwise identities ranging from 80.8% to 100%. The 134 bp insertion elements have homology (>97.0% identity) to both the 5’ and 3’ termini of 5.3 kbp elements annotated in the SG1 genome, the presence of these Long Terminal Repeat (LTR) sequences, as well as the presence and order of domains encoding for *gag* structural polyprotein, aspartic proteinase, integrase, reverse transcriptase and RNAse H, assigned the 5.3 kbp elements to the *Ty1*-*Copia* superfamily of LTR retrotransposons (Wicker et al. 2007). Ten intact, full-length (>5.2 kbp) copies of the transposon were identified in the SG1 genome at a threshold of 95% sequence identity, distributed on chromosomes 5, 6 and 8. Five base pairs at the ends of the 134-bp insertions were directly repetitive on either flank, indicating that they may represent target site duplication sequences (TSDs), which are characteristically 4-6 bp in the case of LTR retrotransposons (Wicker et al. 2007). Promoter analysis determined that all of the 134 bp insertion elements contained an identical putative promoter sequence with a predicted promoter score of 0.99. Those highly-resistant isolates with *PtTi-5* had a second predicted promoter sequence with a score of 0.90 at –213. In the upstream regions of some sensitive isolates (*M2-Ap*) and some highly-resistant isolates (those with *PtTi-*2), there was another predicted promoter sequence at –78 (*M2-Ap*) or –212 (*PtTi-*2), with a score of 0.88. Three additional promoters sequences were predicted in the upstream region of all isolates, at –833 (in wild-types *Sg1-Ap* and *M2-Ap*) or –967 (in *PtTi-1, -2, -3, -4* and *-5*), with a score 0.99, at –498 (in wild-types *Sg1-Ap* and *M2-Ap*) or –632 (in *PtTi-1, -2, -3, -4* and *-5*), with a score 1.00, and at –301 (in wild-types *Sg1-Ap* and *M2-Ap*) or –435 (in *PtTi-1, -2, -3, -4* and *-5*), with a score 0.82. Analysis of transcription factor binding sites within the 134-bp sequences showed that all five elements contained predicted binding sites for ten fungal transcription factors: *ABF1* (2 sites), *ADR1* (1 site), *CBF1* (3 sites), *FACB* (3-4 sites), *GA-BF* (1-2 sites), *GCN4* (1 site), *MAC1* (1 site), *MATα1* (1 site), *NIT1* (1 site) and *RC2* (4-5 sites); all except *PtTi-3* and *PtTi-5* also contained one site for the transcription factor *GAL4* (Figure 2).

### Presence of insertion elements in the *Cyp51A* upstream region is associated with constitutive *Cyp51A* overexpression

Expression of *Cyp51A* was significantly higher in isolates carrying insertion elements in the *Cyp51A* upstream region, both in the presence and in the absence of DMI fungicides, whereas in the wild-type expression was largely induced only in the presence of DMIs (Figure 3).

**Figure 3.**
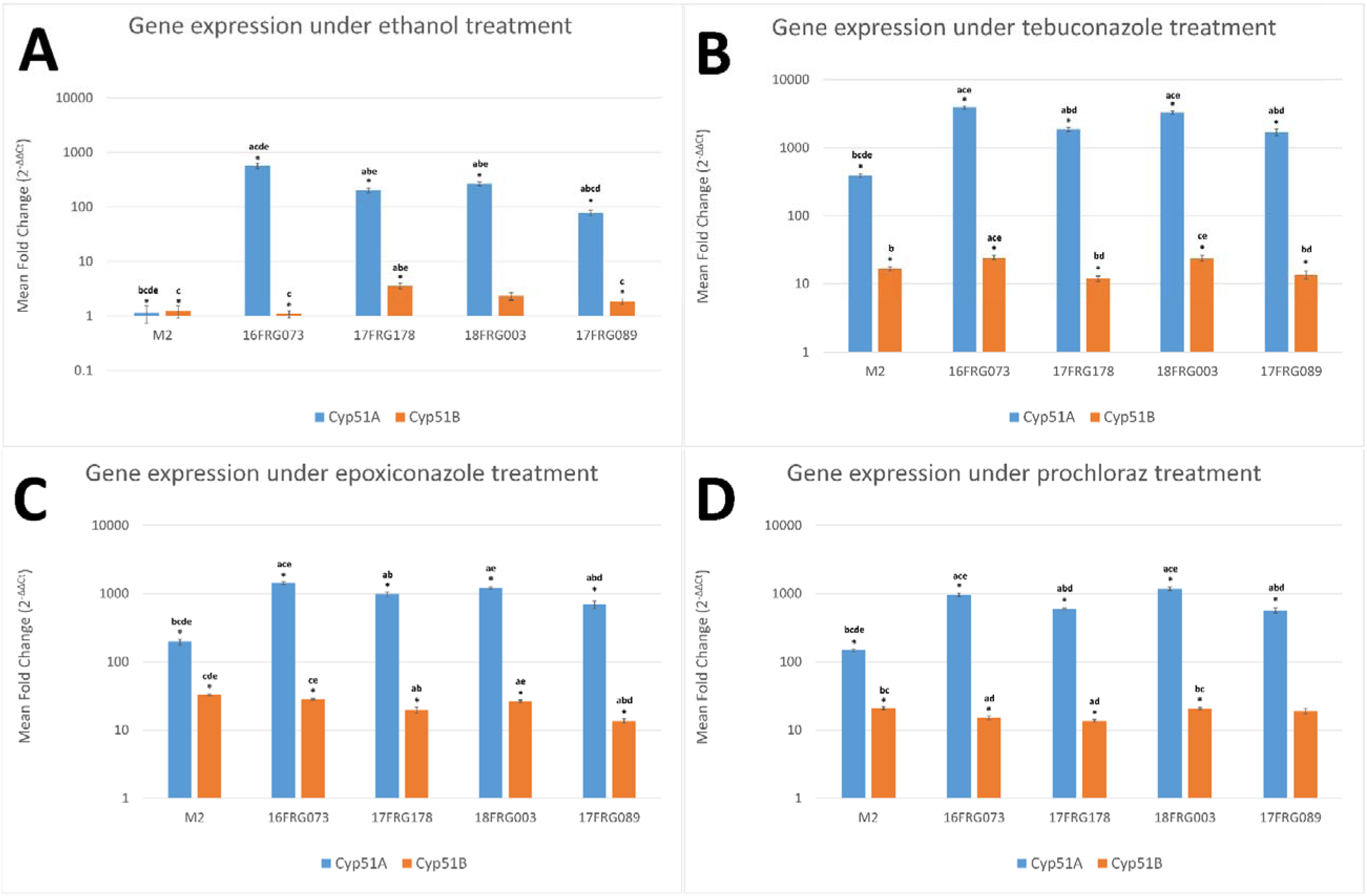
Gene expression of *Cyp51A* and *Cyp51B* in isolates with different variants of the region upstream of *PtmCyp51A*. Isolates M2 (wild-type), 16FRG073 (*PtTi-1*), 17FRG178 (*PtTi-3*), 18FRG003 (*PtTi-4*), and 17FRG089 (*PtTi-2*) were tested under treatment with **(A)** ethanol, **(B)** tebuconazole EC_50_, **(C)** epoxiconazole EC_50_ or **(D)** prochloraz EC_50_. Expression of *Cyp51A* was significantly higher in isolates carrying insertion elements in the *Cyp51A* upstream region, both in the presence and in the absence of DMI fungicides, whereas in the wild-type expression was largely induced only in the presence of DMIs. By contrast, expression of *Cyp51B* was in most cases either lower or not significantly different to the wild-type in isolates carrying insertion elements, in either the presence or absence of DMI fungicides. The results indicate constitutive overexpression of *Cyp51A* (but not *Cyp51B*) in those isolates carrying insertion elements in the *Cyp51A* upstream region, which is further upregulated in the presence of DMI fungicides. Mean fold change of gene expression calculated by 2^−ΔΔCt^ (Livak and Schmittgen 2001), relative to constitutive expression in isolate M2 (ethanol-treated control), normalized to *Actin* as the endogenous control gene (± Standard error of the mean, *n* = 3 biological replicates, four technical replicates per biological replicate). Statistical analysis of 2^−ΔΔCt^ data was performed with IBM SPSS 24.0 using Kruskal-Wallis H test & Dunnett’s T3. *The mean difference between isolates ^a^M2, ^b^16FRG073, ^c^17FRG178, ^d^18FRG003, and ^e^17FRG089 is significant at the 0.05 level.

*Cyp51A* was found to be expressed at very low levels in the ethanol control samples of sensitive isolate M2. Under ethanol treatment in isolates carrying various promoter insertions, expression of *Cyp51A* was 502-fold higher in 16FRG073 (carrying *PtTi-1*), 177-fold higher in 17FRG178 (*PtTi-3*), 234-fold higher in 18FRG003 (*PtTi-4*) and 68-fold higher in 17FRG089 (*PtTi-2*), compared to the wild-type. Kruskal-Wallis analysis showed there were significant differences at the 5% level among the five genotypes tested (*H*(4) = 44.335, *P* < .001). Post hoc tests with Dunnett’s T3 showed the differences were significant (*P* < .005) between each of the genotypes, except for between MR1 isolates 17FRG178 and 18FRG003 (*P* = .256).

Under tebuconazole treatment, expression of *Cyp51A* in DMI sensitive isolate M2 was upregulated 341-fold compared to the same isolate under ethanol treatment. In comparison to the tebuconazole-treated wild-type, in isolates carrying promoter insertions under tebuconazole treatment, *Cyp51A* was expressed 10.1-fold higher in isolate 16FRG073, 4.8-fold higher in 17FRG178, 8.4-fold higher in 18FRG003, and 4.4-fold higher in 17FRG089. Kruskal-Wallis analysis showed there were significant differences at the 5% level among the five genotypes tested (*H*(4) = 51.009, *P* < .001). Post hoc tests with Dunnett’s T3 showed the differences were significant (*P* < .001) between each of the genotypes, except for between 16FRG073 and 18FRG003 (*P* = .138), and between 17FRG089 and 17FRG178 (*P* = .997).

Under epoxiconazole treatment, expression of *Cyp51A* in isolate M2 was upregulated 172-fold compared to the same isolate under ethanol treatment. In comparison to the epoxiconazole-treated wild-type, in isolates carrying promoter insertions under epoxiconazole treatment, *Cyp51A* was expressed 7.2-fold higher in isolate 16FRG073, 4.9-fold higher in 17FRG178, 6.1-fold higher in 18FRG003, and 3.5-fold higher in 17FRG089. Kruskal-Wallis analysis showed there were significant differences at the 5% level among the five genotypes tested (*H*(4) = 43.786, *P* < .001). Post hoc tests with Dunnett’s T3 showed the differences were significant (*P* < .01) between each of the genotypes, except for between 16FRG073 and 18FRG003 (*P* = .130), between 17FRG178 and 18FRG003 (*P* = .259), and between 17FRG178 and 17FRG089 (*P* = .291).

Under prochloraz treatment, expression of *Cyp51A* in isolate M2 was upregulated 128-fold compared to the same isolate under ethanol treatment. In comparison to the prochloraz-treated wild-type, in isolates carrying promoter insertions under prochloraz treatment, *Cyp51A* was expressed 6.5-fold higher in isolate 16FRG073, 4.1-fold higher in 17FRG178, 8.1-fold higher in 18FRG003, and 3.9-fold higher in 17FRG089. Kruskal-Wallis analysis showed there were significant differences at the 5% level among the five genotypes tested (*H*(4) = 49.389, *P* < .001). Post hoc tests with Dunnett’s T3 showed the differences were significant (*P* < .01) between each of the genotypes, except for between 16FRG073 and 18FRG003 (*P* = .098), and between 17FRG178 and 17FRG089 (*P* = .999).

By contrast, expression of *Cyp51B* was in most cases either lower or not significantly different to the wild-type in isolates carrying insertion elements, in either the presence or absence of DMI fungicides. Overall the results indicate constitutive overexpression of *Cyp51A* (but not *Cyp51B*) in those isolates carrying insertion elements in the *Cyp51A* upstream region, which is further upregulated in the presence of DMI fungicides.

In contrast to *Cyp51A, Cyp51B* was found to be expressed constitutively at a relatively higher level in the ethanol control samples of sensitive isolate M2. Under ethanol treatment in isolates carrying various promoter insertions, expression of *Cyp51B* was 0.9-fold lower in 16FRG073, 2.9-fold higher in 17FRG178, 1.9-fold higher in 18FRG003, and 1.5-fold higher in 17FRG089, compared to the wild-type. Kruskal-Wallis analysis showed there were significant differences at the 5% level among the five genotypes tested (*H*(4) = 22.803, *P* < .001). Post hoc tests with Dunnett’s T3 showed the differences between each of the genotypes were not significant at *P* < .05, except for 17FRG178 compared to M2 (*P* = .007), 16FRG073 (*P* = .003) and 17FRG089 (*P* = .045).

Under tebuconazole treatment, expression of *Cyp51B* in isolate M2 was upregulated 13.5-fold compared to the same isolate under ethanol treatment. In comparison to the tebuconazole-treated wild-type, in isolates carrying promoter insertions under tebuconazole treatment, *Cyp51B* was expressed 1.5-fold higher in isolate 16FRG073, 0.7-fold lower in 17FRG178, 1.4-fold higher in 18FRG003, and 0.8-fold lower in 17FRG089. Kruskal-Wallis analysis showed there were significant differences at the 5% level among the five genotypes tested (*H*(4) = 30.533, *P* < .001). Post hoc tests with Dunnett’s T3 showed the differences between the genotypes were significant for 16FRG073 compared to M2 (*P* = .0.12), 17FRG178 *(P* < .001), and 17FRG089 (*P* = .003), and for 18FRG003 compared to 17FRG178 (*P* = .003), and 17FRG089 (*P* = .025).

Under epoxiconazole treatment, expression of *Cyp51B* in isolate M2 was upregulated 26.9-fold compared to the same isolate under ethanol treatment. In comparison to the epoxiconazole-treated wild-type, in isolates carrying promoter insertions under epoxiconazole treatment, *Cyp51B* was expressed 0.8-fold lower in isolate 16FRG073, 0.6-fold lower in 17FRG178, 0.8-fold lower in 18FRG003, and 0.4-fold lower in 17FRG089. Kruskal-Wallis analysis showed there were significant differences at the 5% level among the five genotypes tested (*H*(4) = 36.261, *P* < .001). Post hoc tests with Dunnett’s T3 showed the differences were significant for M2 compared to 17FRG178 (*P* < .001), 18FRG003 (*P* = .014), and 17FRG089 (*P* < .001), for 16FRG073 compared to 17FRG178 (*P* = .018), and 17FRG089 (*P* < .001), and between 18FRG003 and 17FRG089 (*P* < .001).

Under prochloraz treatment, expression of *Cyp51B* in isolate M2 was upregulated 17.1-fold compared to the same isolate under ethanol treatment. In comparison to the prochloraz-treated wild-type, in isolates carrying promoter insertions under prochloraz treatment, *Cyp51B* was expressed 0.7-fold lower in isolate 16FRG073, 0.7-fold lower in 17FRG178, 1.0-fold in 18FRG003, and 0.9-fold lower in 17FRG089. Kruskal-Wallis analysis showed there were significant differences at the 5% level among the five genotypes tested (*H*(4) = 24.575, *P* < .001). Post hoc tests with Dunnett’s T3 showed the differences between each of the genotypes were not significant at *P* < .05, except for M2 compared to 16FRG073 (*P* = .004) and 17FRG178 (*P* < .001), and 18FRG003 compared to 16FRG073 (*P* = .005) and 17FRG178 (*P* < .001).

## DISCUSSION

In this study, 288 *Ptm* isolates collected between 1996 and 2019 were screened for sensitivity to DMI fungicides, and fifty-nine isolates were found to show reduced sensitivity. Four sensitivity levels to DMIs were determined (S, MR1, MR2 and HR) and these were correlated with eight different mutations across the coding sequence and upstream promoter region of the *Cyp51A* gene. The molecular mechanisms underlying the observed phenotypes were categorized as target site modifications due to three different single nucleotide polymorphisms (SNPs) in codon 489, and target overexpression resultant from the insertion of LTR-transposon-derived elements at five different positions in the *Cyp51A* promoter region. Moderately DMI-resistant isolates (MR1 and MR2) had one or the other of these mechanisms whereas highly DMI-resistant isolates (HR) were found to have a combination of both mechanisms together. To our knowledge, this represents the first report of the molecular basis of DMI resistance in SFNB.

The substitution of phenylalanine by leucine at position 489 of the amino acid sequence of the DMI target CYP51A was found in both highly DMI-resistant and select moderately DMI-resistant (MR2) *Ptm* isolates. The same F489L (*F495L*) mutation has been previously reported in Western Australian strains of the closely related pathogen *Ptt*, where it has been associated with reduced sensitivity to DMI fungicides (Mair, Deng, et al. 2016). In *Ptt*, this mutation most affected sensitivity to prochloraz (mean RF 27.7), followed by tebuconazole (mean RF 16.5), metconazole (mean RF 13.8), difenoconazole (mean RF 12.4), propiconazole (mean RF 7.7), prothioconazole (mean RF 2.6) and epoxiconazole (mean RF 1.5) (Mair, Deng, et al. 2016). Structural modelling of different *Ptt* CYP51A isoforms docked with DMI fungicides supported the role of F489L (*F495L*) in altering the catalytic pocket and reducing binding affinities. Of the fungicides tested, the effect of the mutation was greatest on (in declining order) prochloraz, difenoconazole and tebuconazole (Mair, Deng, et al. 2016). The results are broadly in agreement with the patterns of sensitivity found in this study: HR isolates, with CYP51A isoforms differing from those of the *Ptt* mutant isoforms by eight residues (and carrying promoter insertions in addition to F489L), were least sensitive to prochloraz (mean RF 104.9), followed by difenoconazole (mean RF 58.5), tebuconazole (mean RF 50.6), metconazole (mean RF 37.5), propiconazole (mean RF 36.4), prothioconazole (mean RF 23.9) and epoxiconazole (mean RF 9.2). Similarly, the MR2 isolate 18FRG195, with a distinct haplotype incorporating several other unique substitutions (L219F, P225L, I253M) in addition to F489L (*F495L*), showed reduced sensitivity to prochloraz (RF 11.4), metconazole (RF 9.7), difenoconazole (RF 7.0), tebuconazole (RF 6.0), propiconazole (RF 5.8), and prothioconazole (RF 2.7), as well as negative cross-resistance to epoxiconazole (RF 0.5).

In *Ptm*, the phenylalanine to leucine amino acid change was associated with either of three different single nucleotide polymorphisms in codon 489, from the wild-type TTC to either of TTA, TTG or CTC. This suggests that the F489L (*F495L*) mutation has emerged in *Ptm* as a result of three distinct mutational events. This represents a contrast with the situation reported in *Ptt* where all mutant strains had an identical *Cyp51A* sequence and the F489L (*F495L*) substitution was always associated with the same single nucleotide polymorphism, c1467a for codon TTA (Mair, Deng, et al. 2016). A subsequent study using microsatellite markers suggested that this polymorphism most likely arose in *Ptt* from a single mutational event that subsequently recombined into different genetic backgrounds and spread through the pathogen population (Ellwood et al. 2019). By comparison, in this study the mutation F489L (*F495L*) was caused by three different single nucleotide polymorphisms and each of these three SNPs were found in a distinct *Cyp51A* haplotype. This points to multiple independent emergences of the resistant phenotype in the *Ptm* population by a process of convergent or parallel evolution, with the F489 locus representing both an intra-lineage and inter-lineage hotspot of variation (Martin and Orgogozo 2013). Several studies have demonstrated fungicide resistance by a mutation in the target site emerging multiple times independently. For instance, McDonald et al. (2019) has reported eight DMI-resistance mutations in the *Cyp51B* of *Z. tritici* emerging in Australian populations independently of those same mutations originating in European and North American populations, and furthermore the recurrence of particular complex haplotypes of up to seven mutations together were observed in both Australia and Europe. A number of the common mutations in the *Cyp51B* of *Z. tritici*, such as *V136A, I381V* and *S524T*, are also believed to have emerged independently multiple times in European populations (Hawkins et al. 2019). Again in *Z. tritici*, the mutation *G143A* in the *cytochrome b* gene conferring resistance to Quinone outside-Inhibitor fungicides has emerged at least four times independently in Europe (Torriani et al. 2009), and several times independently in North America (Estep et al. 2015). The *G143A* mutation of *cytochrome b* also emerged at least twice independently in European populations of *Plasmopara viticola* (Chen et al. 2007). Multiple independent origins by parallel evolution were also inferred for mutations in the *Sdh* complex genes in Australian populations of *Didymella tanaceti* (Pearce et al. 2019), and for several mutations conferring resistance to different modes of action in North American populations of *Botrytis cinerea* (Fernández-Ortuño et al. 2015). As far as we are aware, the current study represents the only description of independent origins of the same amino acid substitution in the fungicide target site deriving from multiple distinct nucleotide substitutions.

Multiple independent emergence of mutations by parallel evolution also appears to underlay the target overexpression phenotypes found in both highly DMI-resistant and select moderately DMI-resistant (MR1) *Ptm* isolates. Five different 134 base pair insertions were located within the upstream regulatory region of the *Cyp51A* gene and were correlated with constitutive overexpression of the target. Although the insertion sequences shared at least 95% identity, they were found at a number of disparate locations: at positions –46 or –90 to the start codon in highly DMI-resistant isolates, and at positions –66, –74 or –75 to the start codon in certain moderately-DMI resistant (MR1) isolates. Furthermore they appear in a background of two distinct haplotypes of the *Cyp51A* promoter region: those found at –66, – 74, or –75 were in a background that across a 898 bp region upstream was otherwise identical to that of the sensitive reference strain SG1; by contrast those found at –46 or –90 were in a background that was otherwise identical to that of the sensitive strain M2, which differs from SG1 by eight nucleotides across 898 bp. These therefore appear to represent multiple discrete insertion events of *Ty1*/*Copia*-like LTR retrotransposons. This is further reinforced by the five base pair direct repeat sequences at either flank of the LTR-homologous sequences, which are distinct for each of the five insertion elements. These are consistent with Target Site Duplications which are characteristic of transposition events, occurring as a result of staggered double-stranded DNA breaks at the occasion of transposon integration (Linheiro and Bergman 2012), and which characteristically result in 4-6 base pair direct repeats for insertions of LTR transposons (Wicker et al. 2007).

The adaptation of DMI resistance by target overexpression therefore appears to have multiple origins in *Ptm*, with the repeated emergence of insertion mutations within the *Cyp51A* promoter region at different sites and in different genetic backgrounds. This in some respects resembles other cases of multiple insertions in the regulatory regions that have been correlated with *Cyp51* overexpression, such as the four insertion events of 150- to 519-bp elements in the promoter of the drug efflux transporter *MFS1* of *Z. tritici* (Omrane et al. 2017), insertions of 46-, 151- and 233-bp in the *Cyp51B* promoter of *P. brassicae* (Carter et al. 2014), the 1.8-kbp transposon insertion and the 34- and 46-bp tandem repeat elements that have been found in the *Cyp51A* promoter of *A. fumigatus* (Mellado et al. 2007, Albarrag et al. 2011, van der Linden et al. 2013), the 553-bp insertion and the 169-bp tandem repeat element that have been found in the *Cyp51A* promoter of *Venturia inaequalis* (Schnabel and Jones 2001, Villani et al. 2016), and the 126-bp tandem repeat element and the 199-bp insertion in the *Cyp51A* promoter of *P. digitatum* (Hamamoto et al. 2000, Ghosoph et al. 2007). In the most clearly analogous example, Sun et al. (2013) found a 199-bp sequence inserted upstream of both the *Cyp51A* and *Cyp51B* genes in *P. digitatum*, with the insertions sharing a 194-bp consensus sequence deriving from the same Miniature Inverted-repeat Transposable Element (MITE), and differing by their 5-bp target site duplication sequences. In *B. cinerea*, two different rearrangements of the promoter of the *mfsM2* multidrug transporter were both derived from separate insertion events of the same LTR retrotransposon *Boty3* (Mernke et al. 2011). In *Blumeriella jaapii*, four insertion elements of between 2.1- and 5.6-kbp in the *Cyp51B* promoter were derived from at least two separate insertion events of the same long interspersed nuclear element (LINE)-like retrotransposon (Ma et al. 2006). In comparison to the aforementioned cases, where the insertions were generally of different sizes and in most cases different positions within the upstream regulatory region, those described in this study are distinguished principally by their sites of insertion, as they are all of identical size and have a very high degree of sequence similarity between them, apparently deriving from the same or very similar *Ty1*-*Copia* LTR retrotransposons within the *Ptm* genome.

The full *Ty1*/*Copia* LTR transposon to which the 134 bp sequences are homologous is over 5.2 kbp in length in the *Ptm* reference genome SG1, thus the insertion elements found in this study most likely represent Solo-LTRs, which form as a result of the removal of LTR transposons by unequal recombination between similar LTR sequences (Vitte and Bennetzen 2006). This results in the truncation of the internal sequences of the transposon and of one of the Long Terminal Repeats, leaving only a single LTR sequence and the target site duplications remaining at the site of transposon insertion (Zhang et al. 2014, Ji and DeWoody 2016). LTR sequences characteristically contain strong RNA polymerase II promoter elements and transcription factor binding sites, and are capable of independently recruiting transcription factors to drive gene expression (Thompson, Macfarlan, and Lorincz 2016). Solo LTRs are therefore frequently exapted, co-opted as alternative promoters or other *cis*-acting regulatory elements for neighbouring genes (Thompson, Macfarlan, and Lorincz 2016). The LTR-like insertion elements found in this study all contained an identical predicted promoter sequence, as well as a number of predicted transcription factor binding sites. One of these was a predicted binding site for the fungal transcription factor *ADR1*, and interestingly this same transcription factor has been predicted to bind both the 65-bp *Mona* insertion element of *Cyp51B* in *Monilinia fructicola* and the 126-bp tandem repeat element of *Cyp51A* in *P. digitatum* (Hamamoto et al. 2000, Chen et al. 2017). Future promoter deletion and insertion studies may elucidate the core promoter element within the insertion sequences found in *Ptm*, while a Yeast One-Hybrid approach may identify those transcription factors being recruited to drive *Cyp51A* expression. Further investigation is also needed of the activity of the *Ty1*/*Copia* LTR retrotransposons from which the insertion elements are putatively derived. Bioinformatic analysis shows that 10 full-length copies of the transposon are present in the *Ptm* reference genome, and they have normal GC content indicative of low levels of Repeat Induced Point (RIP) mutation activity, implying that these transposons are recently active (Simon Ellwood, personal observation, 01 October 2019). A common feature of retrotransposons across diverse taxa is their activation by biotic and abiotic stresses, with the onset of various adverse living conditions acting as a stimulus to transposon-mediated mutation and genome restructuring events (Zhang et al. 2014, Galindo-González et al. 2017). Furthermore, retrotransposons of the *Ty1*/*Copia* superfamily integrate preferentially to genomic regions that are transcriptionally active (Muszewska, Hoffman-Sommer, and Grynberg 2011). Therefore, it is possible to speculate that the repeated *de novo* emergence of insertions within the *Cyp51A* regulatory region is the result of transposon mobilisation in response to environmental stresses, such as the use of fungicides.

In conclusion, we have found multiple mechanisms which, acting both alone and in concert, contribute to the observed phenomena of DMI fungicide resistance in *Ptm*. Moreover, these mutations have apparently evolved repeatedly and independently in parallel in Western Australian *Ptm* populations. In light of these findings, as well as the ongoing resistance to DMIs in *Ptt*, there is a crucial need for improved fungicide resistance management strategies in the barley net blotch diseases.

---

**Table S1.**
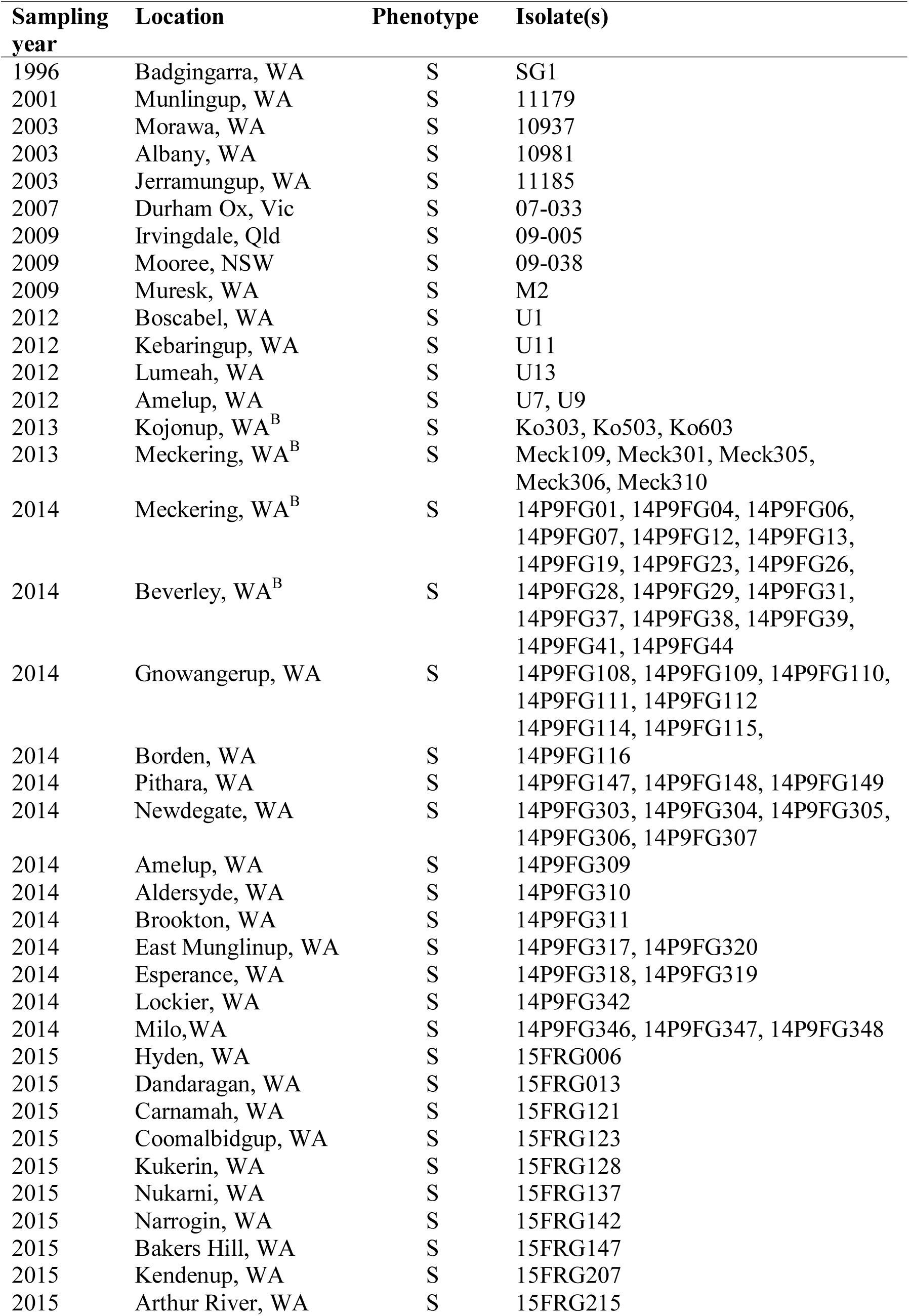
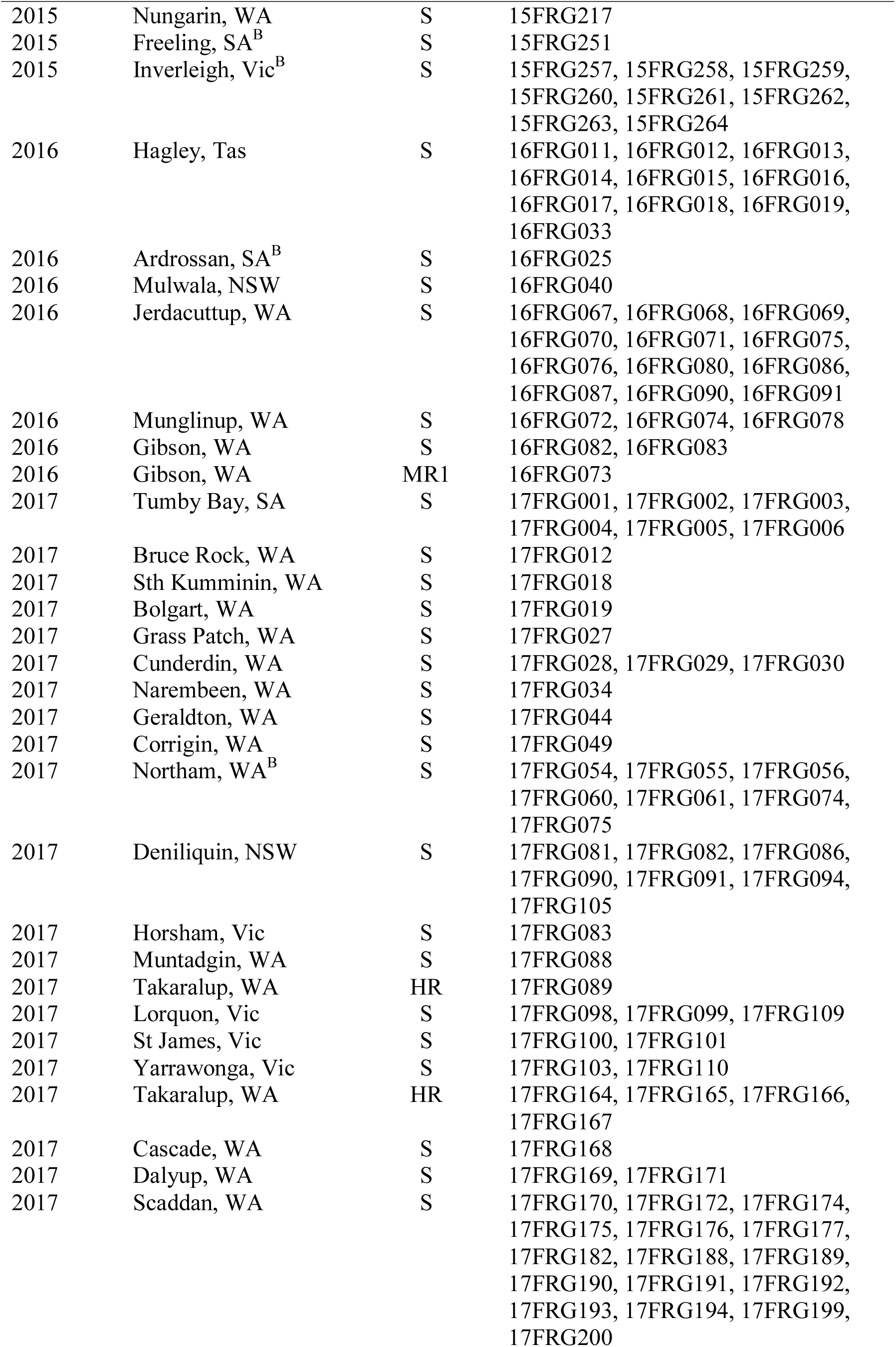
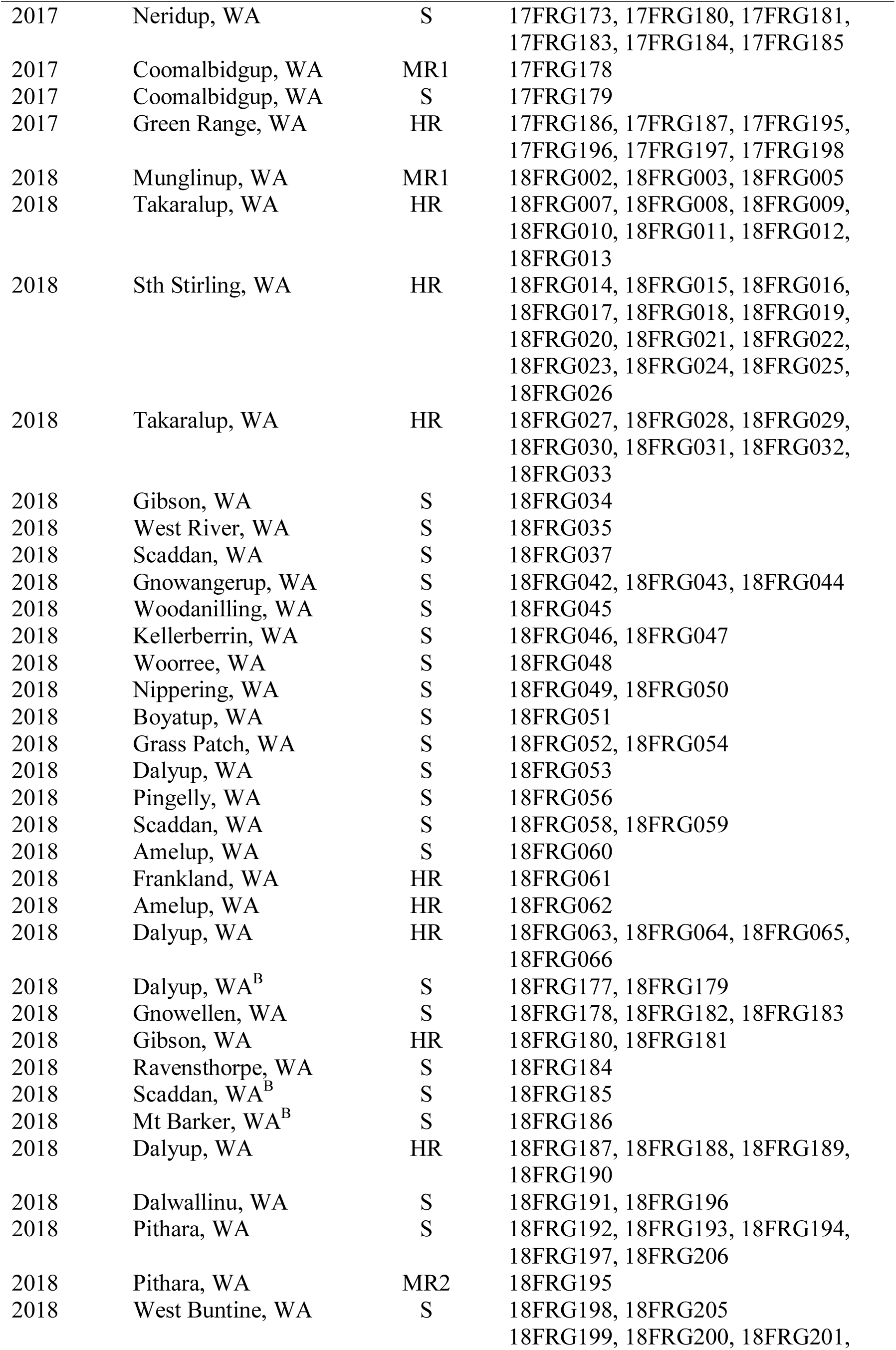
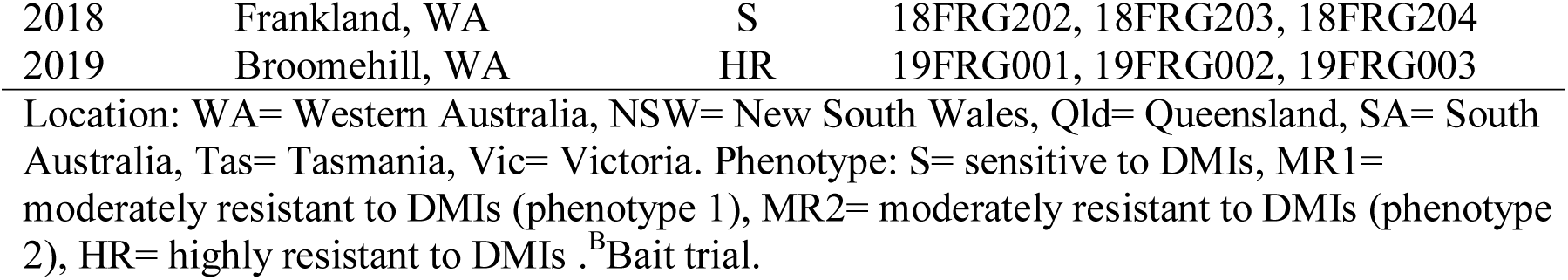
Isolates screened in this study and their relevant characteristics.

**Figure S1.**
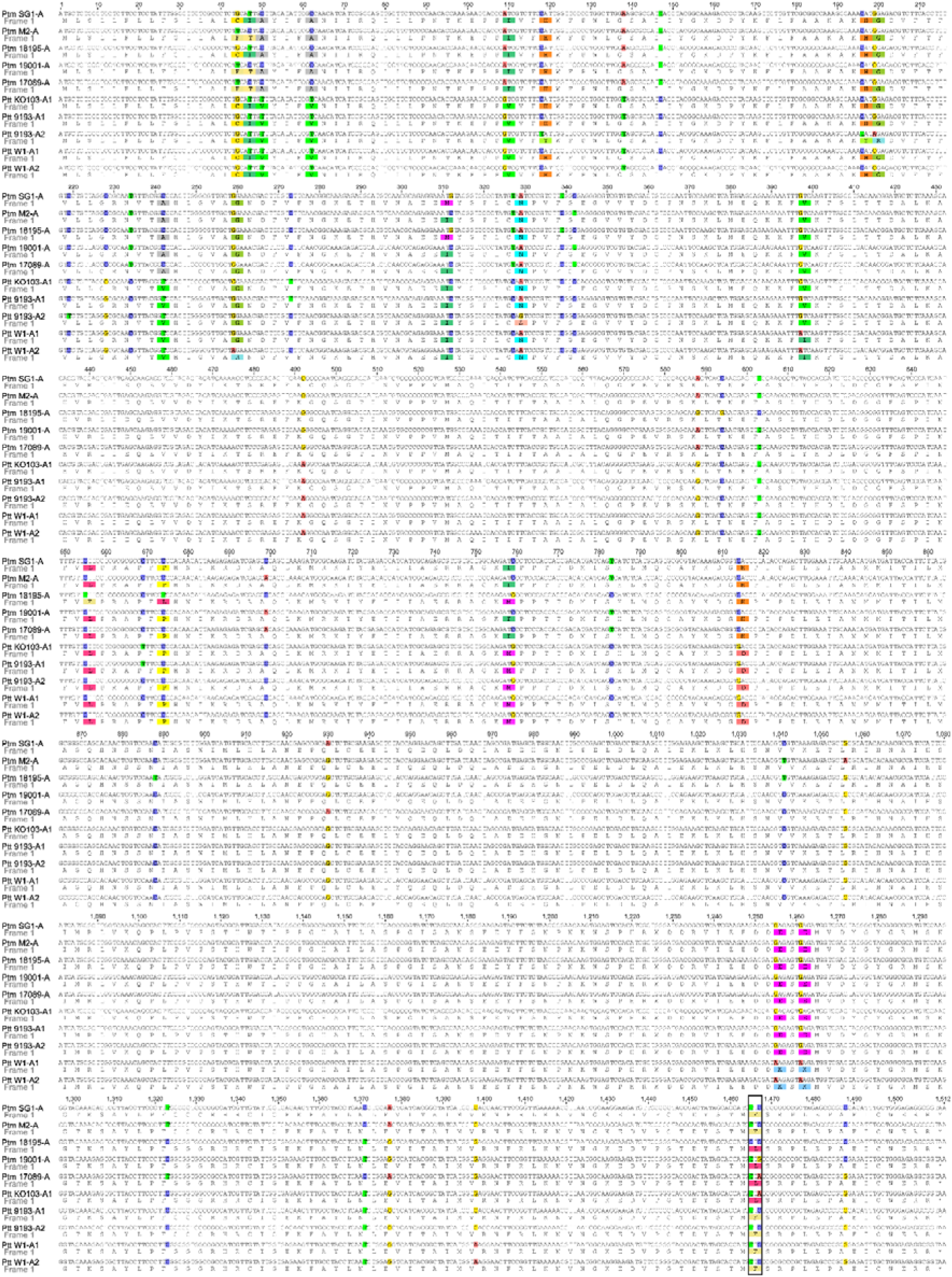
Alignment of complete coding sequences of *Cyp51A* alleles in *Pyrenophora teres* species. Complete 1.512 kbp coding sequences of the five *P. teres* f. sp. *maculata Cyp51A* alleles (*Sg1-A, M2-A, 18195-A, 19001-A, 17089-A*) detected in this study and five *P. teres* f. sp. *teres Cyp51A* alleles (*KO103-A1, 9193-A1, 9193-A2, W1-A1, W1-A2*) reported by Mair, Deng, et al. (2016). Position numbers shown above alignment are relative to start codon in sequence of sensitive reference isolate SG1, translated amino acid sequences are shown below nucleotide sequence, polymorphisms in alignment are highlighted, codon 489 boxed. Alignment generated in Geneious version 6.1.8 software (Biomatters) using ClustalW algorithm with IUB scoring matrix, gap opening penalty 15, gap extension penalty 6.66 and free end gaps.

**Figure S2.**
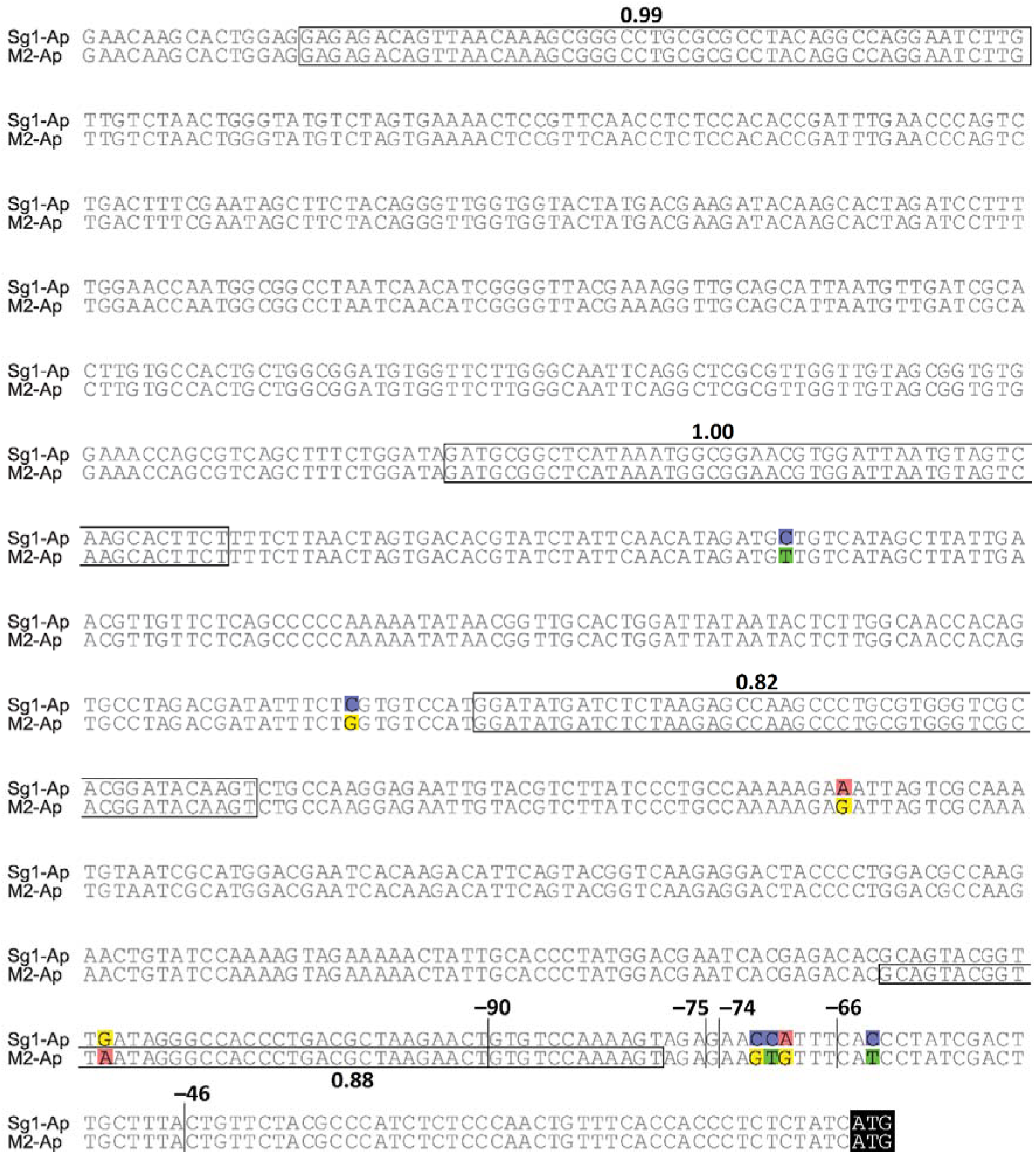
Alignment of upstream region of *PtmCyp51A* in DMI-S isolates. Region 898 bp upstream of the *Cyp51A* start codon in DMI-sensitive isolates carrying alleles *Sg1-Ap* or *M2-Ap*. White box: predicted promoter sequences (predicted promoter scores shown above); Black box: start codons; Vertical bars: sites of insertion of 134 bp elements in MR1 and HR isolates (position numbers shown above are relative to start codon); Polymorphisms in alignment are highlighted in colour. Alignment generated in Geneious version 6.1.8 software (Biomatters) using ClustalW algorithm with IUB scoring matrix, gap opening penalty 15, gap extension penalty 6.66 and free end gaps.

**Figure S3.**
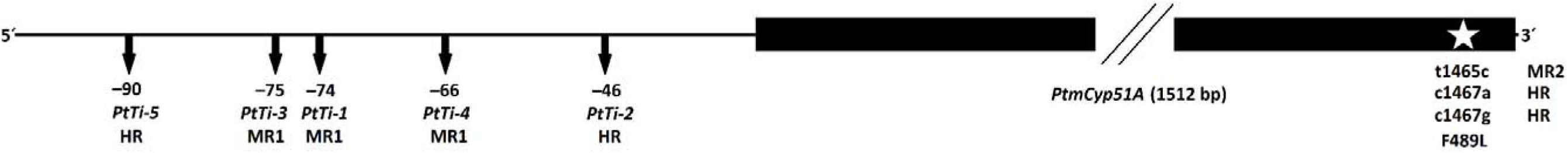
Simplified diagram of *PtmCyp51A* gene and upstream region showing locations of relevant polymorphisms in MR and HR isolates. Position numbers shown are relative to start codon in sequence of sensitive reference isolate SG1. Position of insertion sequences in promoter represented by downwards arrows, position of single nucleotide polymorphisms in coding sequence resulting in substitution F489L in amino acid sequence represented by white star. Insertion sequences of 134-bp in length were found in some moderately-resistant (MR1) isolates at either –66 (*PtTi-4*), –74 (*PtTi-1*), or –75 (*PtTi-3*), and were not associated with any changes in the *Cyp51A* coding sequence. In highly-resistant (HR) isolates, the 134-bp insertion sequences were found at –46 (*PtTi-2*) or –90 (*PtTi-5*), and always in association with the single nucleotide polymorphisms c1467a or c1467g, respectively, both resulting in the amino acid substitution F489L. The amino acid substitution F489L was also found in certain moderately-resistant (MR2) isolates, but resultant from the single nucleotide polymorphism t1465c, and was not associated with an insertion elements in the upstream region.

